# Configurable Compartmentation Enables *In Vitro* Reconstitution of Sustained Synthetic Biology Systems

**DOI:** 10.1101/2022.03.19.484972

**Authors:** Luyao Li, Rong Zhang, Xintong Tian, Ting Li, Bingchun Pu, Conghui Ma, Fang Ba, Chenwei Xiong, Yunfeng Shi, Jian Li, Jay Keasling, Jingwei Zhang, Yifan Liu

## Abstract

The compartmentalized and communicative nature of biological cells contributes to the complexity and endurance of living organisms. Current *in vitro* compartmentalization systems such as droplet emulsions reproduce the compartmentalization property of cells yet fail to recapture the configurability of cellular communication with the environment. To mimic biological cells a step further and expand the capabilities of *in vitro* compartmentalization, we present here a general strategy that inherits the passive transport phenomenon of biology. The strategy incorporates layered, micrometer-sized, hydrogel-based compartments featuring configurability in composition, functionality, and selective permeability of biomolecules. We demonstrated the unique advantage of our strategy in two scenarios of synthetic biology. First, a compartmentalized cell-free protein synthesis system was reconstituted that could support multiple rounds of reactions. Second, we constructed living bacteria-based biosensors in the hydrogel compartments, which could achieve long-lasting functioning with markedly enhanced fitness in complex environments. Looking forward, our strategy should be widely applicable for constructing complex, robust, and sustained *in vitro* synthetic molecular and cellular systems, paving the way for their practical applications.

## Introduction

Compartmentalization is a widespread phenomenon amongst biological systems^1-4^. The existence of physical barriers within and around cells further enables selective retention of essential biological materials while allowing sustained exchange with outer environments. Through compartmentalization, biological cells possess reduced entropy compared to an open system, which contributes to the high efficiency of cellular machinery. It appears that hierarchical compartmentalization enables biological systems to gain increasing complexity and diversity during evolution^5^. For instance, eukaryotic cells envelop genomic materials and gene transcription activities in the nucleus, protecting the genome while regulating key cellular events at levels unavailable to prokaryotes^6,7-10^. Moreover, many compartments of cells are dynamic and configurable. The permeability of bacteria cells can be tuned to resist environmental impacts through sophisticated mechanisms^11-13^. The configurable compartmentalization of cells at multiple hierarchical levels contributes to biological organisms’ richness and diversity.

Inspired by nature, we developed a <u>con</u>figurable <u>c</u>ompartmentation strategy for <u>e</u>ngineered *in vitro* molecular and cellular systems (ConCEiV). Using a microfluidic approach, ConCEiV encapsulates these systems in micrometer-sized, layered hydrogel compartments. Hydrogels are ideal materials for accommodating biomolecules and living cells. They provide a semi-permeable aqueous environment, allowing sustained reagent exchange for diffusion-limited biochemical reactions and cell culture and functioning. The configurability of our hydrogel-based compartments lies in the ease of tuning composition and geometry, functionalization (e.g., magnetizing), and, importantly, deterministic adjustment of selective permeability (Fig. 1). These features endow ConCEiV with broad applicability of reconstituting *in vitro* systems across different hierarchies. We demonstrate in this work the compartmentalization of a cell-free protein expression system for sustained, multi-cycle production of target proteins, and a higher-hierarchy system – an engineered living bacteria-based biosensor exhibiting enhanced environmental fitness.

**Figure 1.**
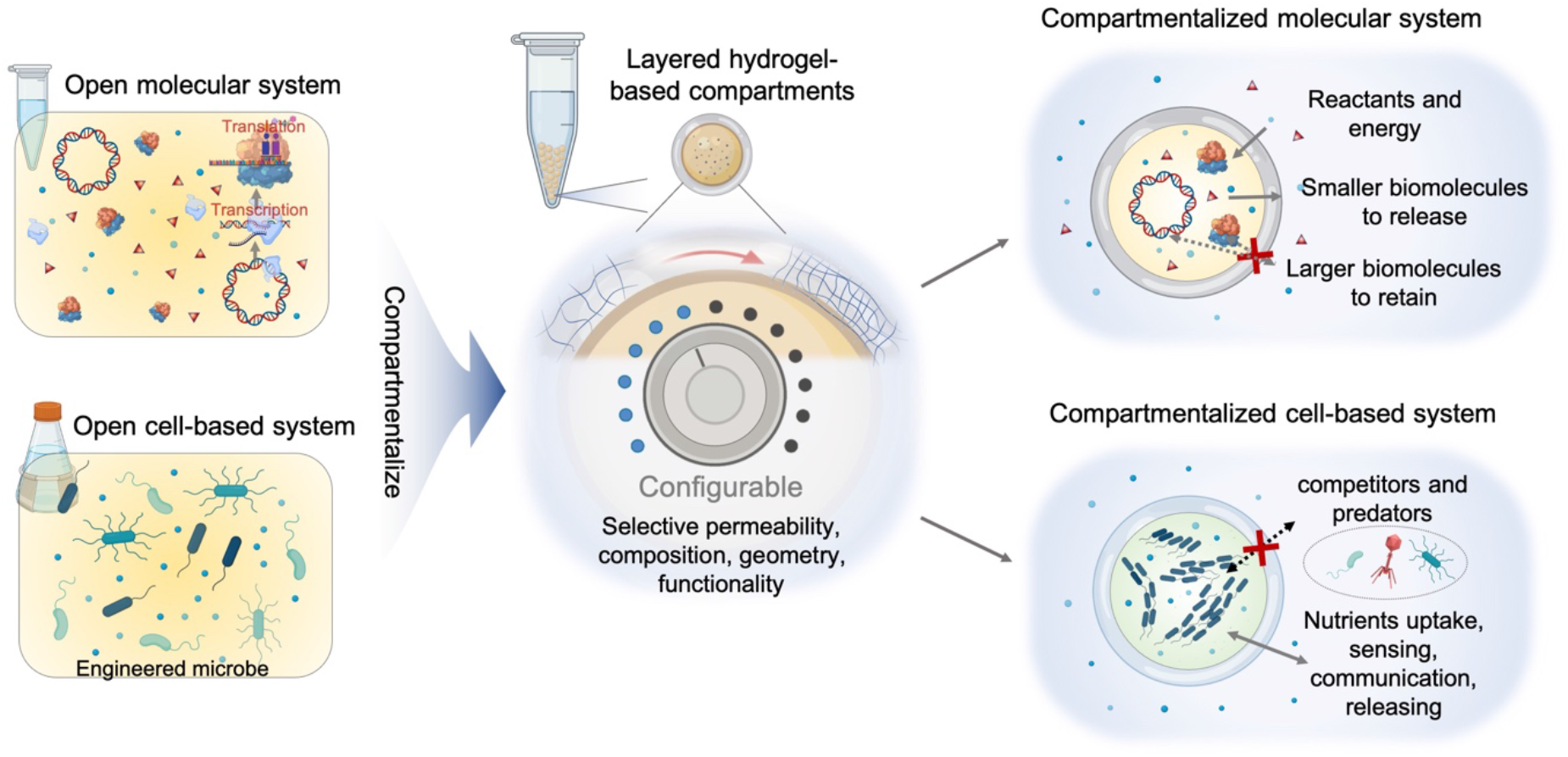
Schematic of the ConCEiV strategy. ConCEiV compartmentalizes open *in vitro* molecular and cell-based systems in micrometer-sized, layered hydrogel-based compartments. The compartments own configurability in selective permeability, composition, geometry, and functionalization. The compartmentalized systems allow biological reactions and activities to execute in a sustained manner due to configured passive transport. For a reconstituted molecular system, the selective permeability of compartments can be tuned to achieve desired transport of biomolecules such as protein and DNA while allowing constant input of reactants and energy. Reconstituted cell-based systems can protect the encapsulated engineered cells from competitors and predators and support the cells’ growth and functioning, thereby enhancing the fitness of these functional synthetic biology systems in complex environments.

Over the years, efforts have been devoted to reconstituting cell-like compartments *in vitro*, aiming to study the complexity of cells from the bottom-up^14-16^, and to perform highly parallel and heterogeneous reactions in a discrete manner^17, 18^. A typical *in vitro* compartmentalization strategy is water-in-oil (w/o) emulsification which yields isolated aqueous compartments surrounded by an immiscible oil^19^. Coupled *in vitro* transcription and translation within these w/o droplets have been realized by Tawfik and Griffiths to select desired genes^20^. Although micelle- or vesicle-based transport does allow the exchange of small organic molecules across surfactant-stabilized w/o droplets, many bioactive molecules are hydrophilic, and the continuous oil phase prevents molecular transport between compartments and their surroundings^21^. Hence, it is highly desirable to build a sustained and interactive *in vitro* system which appreciates an aqueous environment with selectively permeable compartmentalization. Such a system could be living material-based biosensors to monitor environmental signals or bioreactors fueled by external energy/nutrients. Compartmentalization can be realized in an all-aqueous setting by creating membrane-based vesicles such as liposome^22^, proteinosome^23^, and polymersome^24^. Some of these membranes are intrinsically porous, or nanopores can be intentionally created to acquire selective permeability^25, 26^. These membrane-based compartments are inherently fragile, limiting their practical adoption in field studies and real-world environments. Moreover, engineering these compartments to meet highly specified and customized permeability requirement remains challenging. With our layered hydrogel-based compartmentalization, ConCEiV provides an elegant solution to these challenges and offers new desired capabilities in reconstituting *in vitro* systems.

Synthetic biology is a multidisciplinary area of research that seeks to create new biological parts, devices, and systems^27, 28^. Cell-free protein synthesis (CFPS) is an emerging field in synthetic biology which enables inexpensive and fine control of recombinant protein expression^29, 30^. In CFPS, the physical boundary of cells, the cell membrane, is disrupted to form an open *in vitro* environment, rendering the flexibility for adding or removing natural/synthetic components. With ConCEiV, we re-compartmentalize CFPS and artificially tune the selective permeability of compartments to achieve spatially decoupled *in vitro* transcription and translation (TX-TL): the plasmid DNA template and RNA polymerase (RNAP) are confined within the gel membrane, while transcribed messenger RNAs (mRNAs) transport freely to the external buffered solution, where they continue to be translated for producing target proteins. In addition to the pieces of machinery required for translation, the buffered solution also contains materials and energy for transcription, which continuously diffuse into the compartments through the hydrogel membrane. The decoupled TX-TL allows us to preserve transcription within the microgels by recycling plasmids and RNAP, and replenish translation by feeding fresh solutions, thereby enabling multi-cycled, sustained CFPS.

On the other hand, building higher hierarchical systems based on engineered living cells can open up a wealth of new applications^27^. The extensive portfolio of synthetic biology toolkits offers myriads of genetically modified microorganisms (GMMs) with genetic circuits enabling living diagnostics and therapeutics^31^, programmable living materials^32^, information storage^33^, and environmental sensing^34^. However, these functional genetic circuits exert an extra metabolic burden and decrease the GMM’s fitness in a highly complex natural environment^35, 36^. How may we protect these engineered strains against predators like bacteriophage or their fellow microbes^37^? Furthermore, after they perform each intended task, it would be highly desirable to retrieve these GMMs to extract their recorded information. While considerable attention has been paid to contain GMMs from escaping to natural environments^38^, protecting GMMs in real-world settings for sustained functioning is currently neglected. A competent and deployable protection strategy that improves the fitness of GMMs is yet to be developed. ConCEiV provides a feasible solution to this challenge by compartmentalizing GMMs in layered microgels. The core hydrogel houses the cells and provides a cozy environment for cellular growth and function. The shell hydrogel is designed to have a dense polymer network that prevents external microbial competitors and predators from invading while simultaneously allowing the exchange of nutrition and molecular signals. Bacteria-based living sensors prepared by ConCEiV show near-perfect resistance against wild bacteriophages and maintained functionality. We further demonstrate that ConCEiV enables easy retrieval of living sensors after monitoring pollutant levels in synthetic river silt samples and subsequent high-throughput analysis to retrieve recorded pollutant information by flow cytometry.

## Results

### Manufacturing of ConCEiV compartments

The ConCEiV strategy compartmentalizes *in vitro* systems within layered microgels through a two-step droplet microfluidic approach (Fig. 2a). First, engineered molecular or cellular species are co-encapsulated with hydrogel solution using a flow-focusing microdevice (Supplementary Fig. 1a). The generated droplets are then gelled and de-emulsified, yielding monodispersed gel cores. Second, the core beads are reinjected into a dual-junction flow-focusing device (Supplementary Fig. 1b) to build the shell layer. Due to the close-packed nature^39^ of gel beads in microchannels, core-shell hydrogel droplets can be generated at ∼90% efficiency (Supplementary Fig. 2). Gelling and demulsification are repeated to obtain the final layered hydrogel compartments. The entire procedure can be completed in 3 hours, yielding over 3,000,000 compartments.

**Figure 2.**
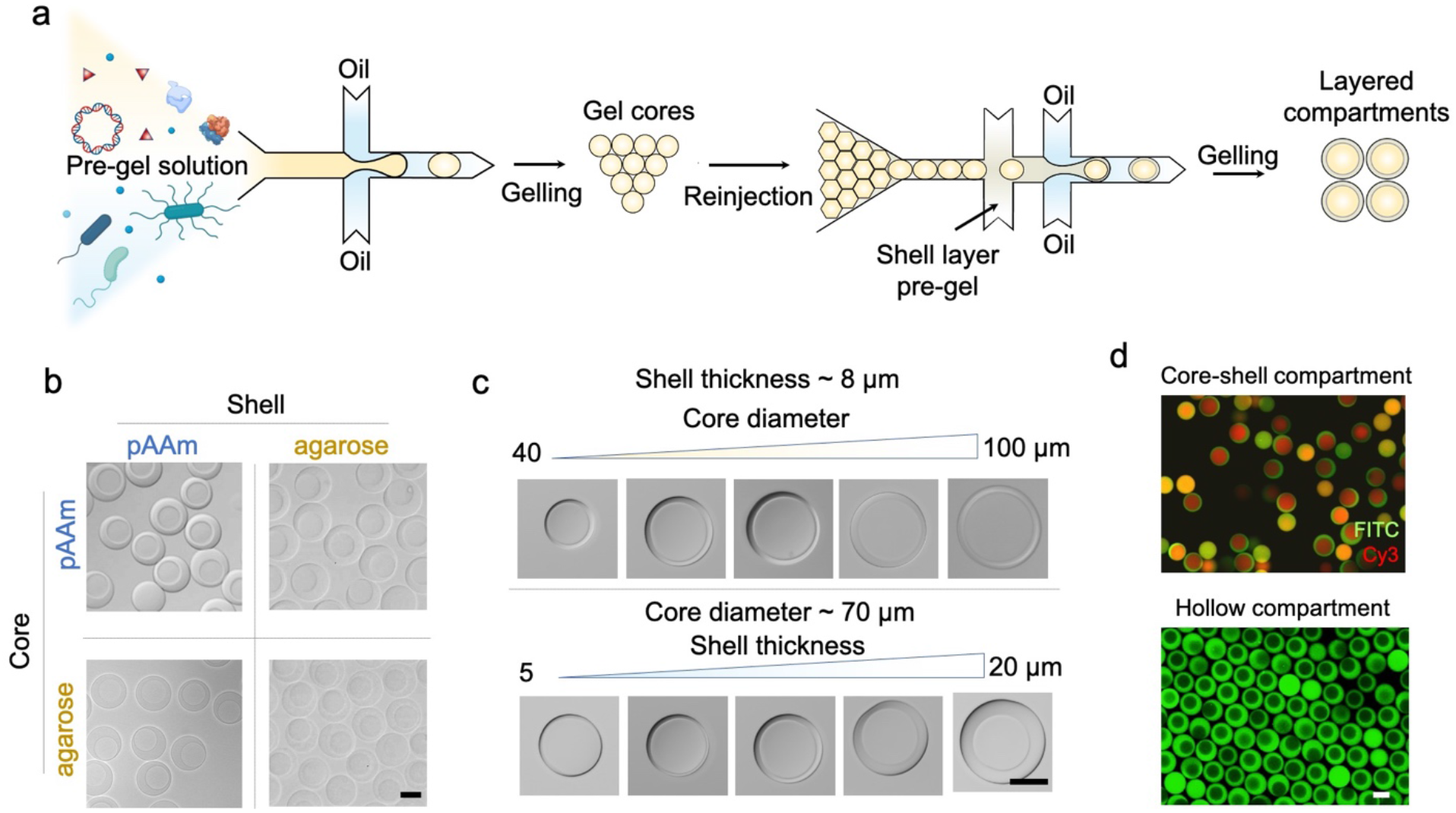
Manufacturing of the ConCEiV compartments. (a) Microfluidic workflow to encapsulate engineered synthetic biology systems in layered compartments. The systems are pre-encapsulated in the hydrogel cores using a microfluidic dropmaker. The core beads are further reinjected to a second microfluidic device to generate layered compartments. (b-d) Optical micrographs of (b) compartments composed of arbitrary combinations of polyacrylamide (PAAm) and agarose hydrogels, (c) compartments featuring tunable core diameters between 40 to 100 μm and shell thickness between 5 to 20 μm, and (d) core-shell hydrogel compartments and hollow compartments after core dissolving. The core and shell layer of compartments in (d) are fluorescently labeled with Cy3 (red) and FITC-dextran (green), respectively. Scale bars: 50 μm.

### Configuring composition and geometry of compartments

The two-step workflow of ConCEiV offers the ease of altering both core and shell hydrogel materials on demand. Fig. 2b shows manufactured core-shell compartments composed of arbitrary combinations of polyacrylamide (PAAm) and agarose hydrogels. PAAm gel typically features a dense polymer network favorable for confining biomolecules. Agarose gel features mild radical-free polymerization, thus being more friendly to accommodate living cells. Moreover, the workflow allows precise tuning of the compartment geometry (Fig. 2c). The core diameter and shell thickness are adjustable between 40 – 100 and 5 – 20 μm, respectively, by simply modifying the microfluidics. If desired, the core of compartments can be selectively dissolved to create an all-liquid cavity suitable for biological reactions. Fig. 2d displays the manufactured compartments featuring a core-shell architecture with each layer labeled by distinct fluorescence. The core gel is reversibly crosslinked, which can be completely dissolved within 3 min (Supplementary Fig. 3&4), leading to a hollow architecture. The compartments can also be conveniently magnetized by involving ferric oxide nanoparticles, thereby enabling efficient recycling and buffer exchange (Supplementary Fig. 5).

### Configuring selective permeability of compartments

The selective permeability of hydrogels has been extensively utilized to sieve biomolecules^40^. We hypothesized that the hydrogel shell of ConCEiV compartments could serve as a similar molecular sieve, selectively barring the transport of biomolecules on our demand. To test our hypothesis, we designed a ConCEiV-based polymerase chain reaction (PCR). As illustrated in Fig. 3a, DNA plasmids are pre-encapsulated in the liquid cavities of compartments as PCR templates and assumed to be physically confined therein. A PCR mix is externally introduced and infused into the compartments. Suppose the physical dimension of amplicon DNA (double of its gyration radius *R*_*g*_) is greater than the pore size of hydrogel shell (*D*_*p*_). In that case, they are expected to accumulate in the cavity during the amplification, whereas smaller products are rather free to transport to the outer environment. A preliminary experiment was first conducted, confirming that our customized PCR strategy was conducted as designed (Supplementary Fig. 7). We then manufactured a set of compartments with three distinct pore sizes of the shell. This was accomplished by tuning the ratio of crosslinker, bis-acrylamide (Bis), of a PAAm gel. Determined by scanning electron microscopy, the average *D*_*p*_ of a PAAm gel containing 0.6%, 0.45%, and 0.24% Bis is 31, 58, and 131 nm, respectively (Supplementary Fig. 6). We next designed ten primer pairs targeting amplicons ranging from 150 to 1187 bp on the plasmid and performed a series of ConCEiV-based PCRs (Supplementary Fig. 8). Fig. 3b displays the fluorescence distribution of compartments (Supplementary Fig. 9) subject to various PCR conditions and a set of micrographs showing representative 0.6% Bis-PAAm compartments post PCR. Overall, the DNA amplicons undergo a mobility transition from being diffusible through the shell to being confined in the cavity with the increase of their length, which can be noted from the elevated fluorescence profile. For each different *D*_*p*_, however, the transition occurs at distinct stages; the threshold length at which most of the amplicons can be trapped is estimated to be ∼ 434, 547, and 866 bp for *D*_*p*_ of 31, 58, and 131 nm, respectively, showing a positive correlation with *D*_*p*_. These results verify that our compartments indeed possess a size selection effect of DNA molecules, and, importantly, the selective permeability can be readily configured by simple adjustment of the hydrogel shell composition.

**Figure 3.**
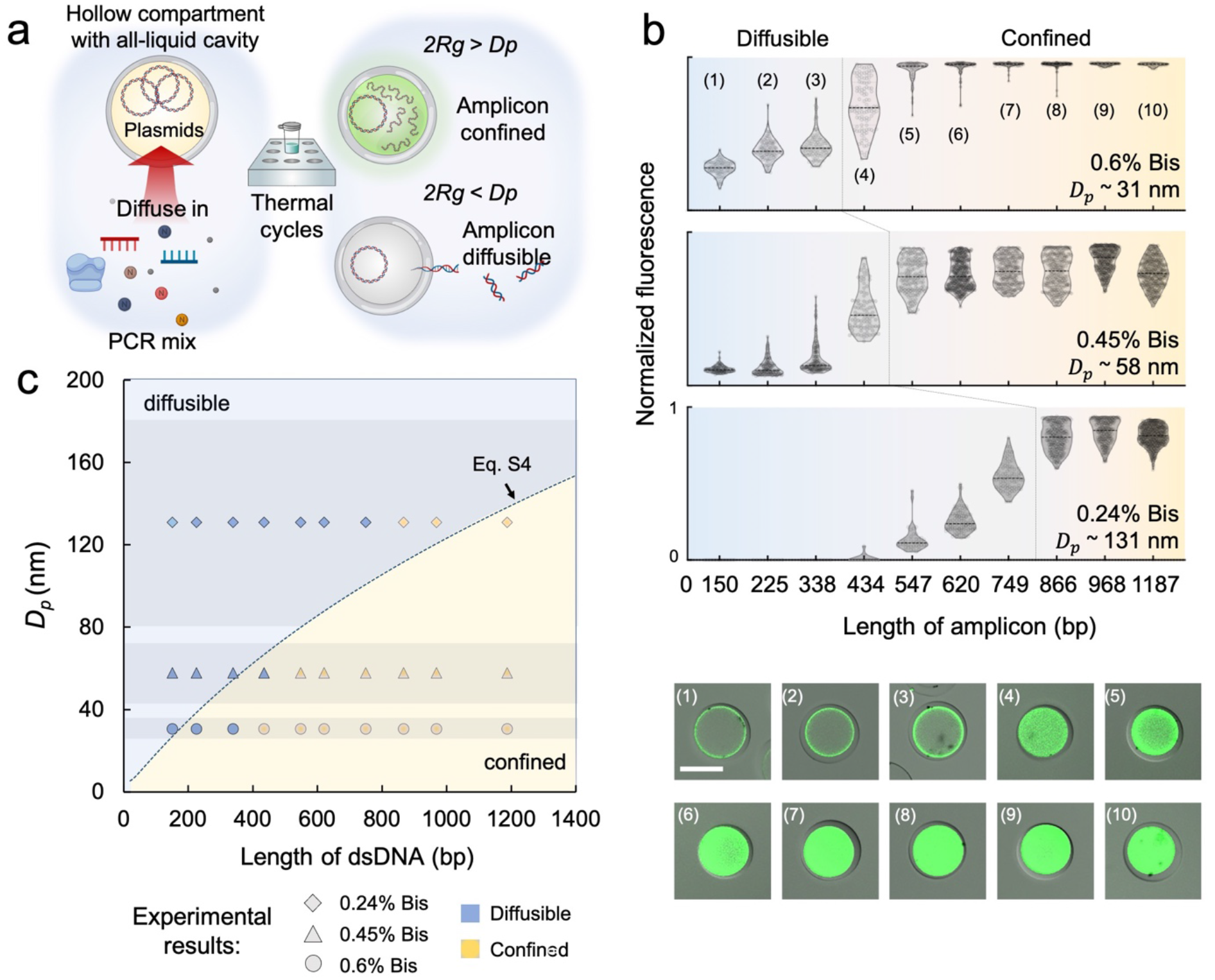
Configuring selective permeability of the ConCEiV compartments. (a) Schematic of a compartmentalized PCR assay designed to explore the selective permeability of compartments. If the dimension of amplicon DNA (double of its gyration radius *R*_*g*_) is greater than the pore size of compartment shell (*D*_*p*_), they are expected to be accumulated in the cavity. By contrast, smaller amplicons will diffuse to the outer solution. The process can be visualized by fluorescent labeling of the amplicons. (b) Results of compartmentalized PCR performed in compartments with configured pore sizes. The scatter plots (top) display fluorescence profile (normalized) of the compartments subject to serial PCRs. Each point in the plots represents the value of a single compartment. The series of fluorescent micrographs (bottom) show individual 0.6% Bis-PAAm compartments undergone corresponding PCR conditions. Scale bar: 50 μm. (c) Physical model describing the transport characteristic of dsDNA across the compartments. The scattered points represent experimental results (legend) obtained from the compartmentalized PCR. The shadowed areas represent the extent of standard deviation of *D*_*p*_ calculated from SEM measurements (Supplementary Fig. 6).

It would be beneficial if we could predict the selective permeability of our compartments for any given pore size without performing the customized PCRs. With this goal, we built a physical model considering the free diffusion of semiflexible DNA molecules in porous media (Supplementary Materials). The model approximates that the diffusible-to-confined transition takes place where the geometrical size of DNA is close to the size of hydrogel pores. In an aqueous solution, the geometry of DNA as a semiflexible polymer can be characterized by its radius of gyration using the Kratky-Porod equation^41^ (Supplementary Materials, Eq. S1). Based on this, we derived the relationship between the critical transition length of DNA and *D*_*p*_ (Supplementary Materials, Eq. S4). Fig. 3c plots model-defined diffusible and confined regimes divided by Eq. S4. For comparison, we included the experimental data derived from Fig. 3b. As can be seen, the model exhibits reasonable agreement with the experimental results. Therefore, we believe that the established model can provide a preliminary guide for designing selective permeability of our compartments to confine or release desired DNA molecules.

### Establishing ConCEiV-based cell-free protein synthesis

Having examined the configurability of the compartments, we next sought to reconstitute compartmentalized protein synthesis *in vitro* toward a sustained, long-lasting system. Typical CFPS assays are batch reactions where synthesized protein products and the molecular machinery are mixed together^42^. The latter are not recyclable and often discarded in subsequent protein purification steps, limiting the total cost-effectiveness and disobeying the central goal of synthetic biology for green biosynthesis. With ConCEiV, we aimed to bridge compartmentalized biological *in vivo* protein expression and open-environment CFPS, creating a semi-open protein synthesis system *in vitro*. As shown in Fig. 4a, ConCEiV-based protein synthesis consists of hollow compartments with their liquid cavities containing genes and enzymes immersed in a feeding solution. We hypothesized that by configuring the selective permeability, the plasmid templates and RNAP could be retained within the compartments, allowing for RNA transcription. Other required materials for transcription, such as dNTPs and energy, can continuously diffuse in from the outer feeding solution. The transcribed mRNA could then transport back to the feeding solution that contains the translation machinery to initiate protein synthesis.

**Figure 4.**
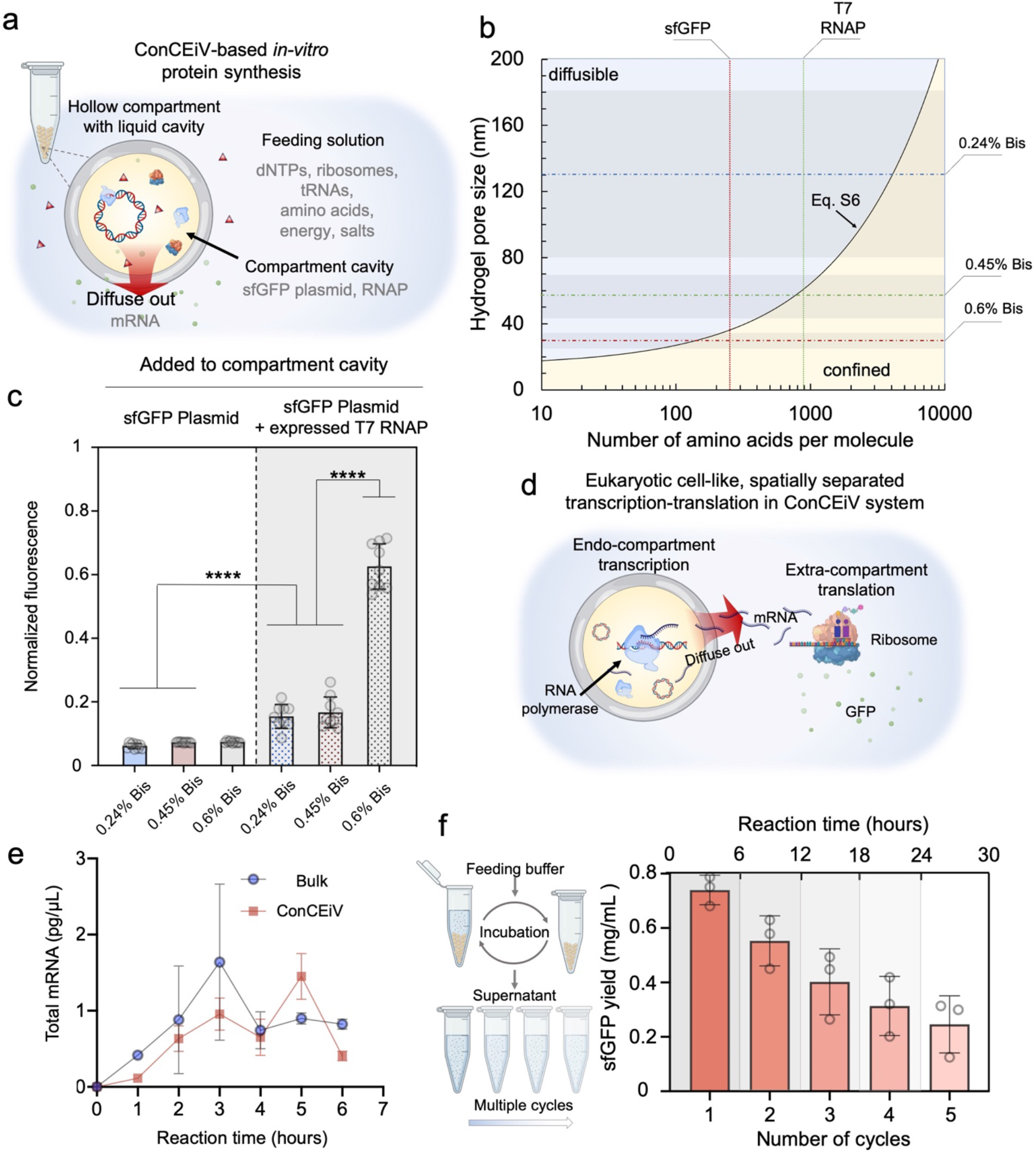
ConCEiV-based *in vitro* protein synthesis. (a) Schematic of the compartmentalized protein synthesis system. Hollow Bis-PAAm compartments encapsulating plasmid and RNA polymerase (RNAP) are immersed in a feeding solution containing other required reagents for RNA transcription and protein translation. (b) Physical model describing the transport characteristic of globular proteins across the compartments. The dashdotted lines represent the averaged pore size of three different Bis-PAAm gels with corresponding shadowed areas denoting the extent of the standard deviation. Vertical dotted lines indicate total amino acid number of the stated protein molecules. (c) sfGFP fluorescence levels (normalized) of protein synthesis assays performed on three different Bis-PAAm compartments. In the 1^st^ group (bright background) of experiments, solely sfGFP plasmid was added to the compartment cavity. *E. coli* cell extract containing *in vivo* expressed T7 RNAP was added along with sfGFP plasmid in the 2^nd^ group (grey background). Error bars denote standard deviations of multiple parallel experiments (n ≥ 8). Statistical analysis: unpaired two-tailed t-test. (d) Proposed transcription-translation procedure in the ConCEiV-based system. Transcription and translation are spatially separated. Plasmids and RNAP are confined in the compartments for transcription. The produced mRNAs transport to the feeding solution where translation takes place. (e) Total mRNA levels during sfGFP expression by a ConCEiV-based system and a bulk CFPS system. Error bars denote standard deviations of multiple parallel experiments (n = 3). (f) Workflow (left) and yield (right) of multicycle sfGFP synthesis by a ConCEiV-based system. For each cycle the 0.6% Bis-PAAm compartments containing sfGFP plasmid and cell extract were incubated in a fresh feeding solution for 6 hours, after which compartments were magnetically recycled and the sfGFP yield in the supernatant was quantified.

With the established DNA transport model (Fig. 3c), we were able to expect that a 2.48 kb plasmid encoding superfolder green fluorescent protein (sfGFP) would be confined in the compartments. Even taking supercoiling into account does the physical size of the plasmid exceed the largest pore size in our case^43^. To predict the configurability of protein transport across the compartments, we modified the DNA model by replacing the gyration radius of DNA with that of a globular protein molecule given its total number of amino acids, which yielded a brief model describing the diffusible-to-confined characteristics of proteins (Fig. 4b, Supplementary Materials, Eq. S6). Again, we marked the pore size profile of 0.24%, 0.45%, and 0.6% Bis-PAAm hydrogel shell on the plot. The model suggests that T7 RNAP, comprising 883 amino acids, would be confined in 0.6% Bis-PAAm compartments with an average pore size of 31 nm. By contrast, sfGFP, having only 241 amino acids, could transport through the compartments with the marked pore sizes. It is noteworthy that bacteria ribosome is a vast RNA-protein complex, with the number of amino acids exceeding 7,000^44^. Thus, we expect that ribosomes could not transport through all of these compartments.

With insights gained from the models, we moved to constitute ConCEiV-based *in vitro* protein synthesis systems experimentally. We first encapsulated only sfGFP plasmids in the compartments and incubated them in an *E. coli* extract-based CFPS mixture. The sfGFP fluorescence is weak for all shell pore size conditions (Fig. 4c), suggesting that the TX-TL procedure could barely occur. This is possibly due to the hydrogel shell’s physical barrier effect, which limits the effective transport of RNAP into the compartment cavity to initiate the transcription. Our model suggests that RNAP could transport into the 0.24% and 0.45% Bis-PAAm compartments, yet the limited local concentration might lead to extremely low transcription efficiency. Further, we added crude cell extracts containing *in vivo* expressed T7 RNAP to the compartment. As a result, we observed notable increment in sfGFP fluorescence, suggesting that the transcription events indeed occurred in the cavity. Most importantly, the fluorescence level of 0.6% Bis-PAAm compartments is significantly elevated, 8.5 folds of the previous value where only plasmid was added and over 3.5 folds of those compartments with larger pores (0.24%-0.45% Bis). This reveals a much more efficient transcription occurring within 0.6% Bis-PAAm compartments, which suggests that 0.6% Bis-PAAm compartments enable a significantly higher confinement effect on RNAP than that of other compartments, in substantial agreement with our model prediction. We further derived that the sfGFP yield for 0.6% Bis-PAAm compartments was as high as 0.74 mg/mL, comparable to that of recently reported open CFPS systems^45^. To confirm the effect of confined RNAP on protein expression, we added an extra dose of purified RNAP to 0.6% Bis-PAAm compartments and observed a further increased sfGFP yield (Supplementary Fig. 12). This also suggests that the protein yield of the ConCEiV-based CFPS system can be further improved upon additional optimization^42^. These findings verified the successful establishment of a ConCEiV-based *in vitro* protein synthesis system. As illustrated in Fig. 4d, the system separated transcription and translation in space by confining plasmids and RNAP in the compartments while allowing the transport of mRNA to the outer feeding solution, thus achieving a eukaryotic cell-like, spatially decoupled TX-TL process. To further verify this, we monitored the mRNA level in the feeding solution throughout the protein synthesis procedure (Fig. 4e). The mRNA level of our ConCEiV-based system is comparable to that of bulk CFPS, confirming the reasonably high efficiency of the endo-compartment transcription and efficient transport of the mRNAs to the outer solution.

### Multicycle *in vitro* protein synthesis enabled by ConCEiV

Having established an optimal ConCEiV-based protein synthesis system, we wanted to test whether our proposed goal – sustained and multicycle protein synthesis could be achieved. To do this, we encapsulated sfGFP plasmids and RNAP in 0.6% Bis-PAAm compartments, which were subsequently immersed in the feeding solution for protein synthesis. After each cycle, the feeding solution was collected, and the compartments were magnetically recycled to incubate with new feeding solutions. We repeated five cycles and quantified the sfGFP yield of each cycle (Fig. 4f). Although a decay of the expression level was observed along with the process, possibly due to the deactivation of RNAP after prolonged functioning, a yield of 0.25 mg/mL was still obtained at the fifth cycle. The overall yield of the 5-cycled expression exceeded 2.2 mg/mL, around 3 folds of its first cycle. These results verified that we have successfully achieved multicycle *in vitro* protein synthesis, a key step moving toward green and sustainable biosynthesis.

### Establishing ConCEiV-based living biosensors

We next attempted to compartmentalize a higher-hierarchy system – living bacteria-based biosensors. We envisioned that the ConCEiV system could serve to encapsulate, protect and retrieve GMMs, enabling robust and sustained functioning in complex environments. As a demonstration, we explored the ConCEiV architecture to grow a valerolactam biosensing strain in a core hydrogel matrix. At the same time, a tougher semi-permeable outer layer served as a barrier against outside predators, such as phages (Fig. 5a). Lactam is an essential class of Nylon precursor and could be detected in the environment as an industrial chemical pollutant^46^. The concentration of lactam residual in the waste falls within 100 mM range^47^. To build lactam-responsive biosensors, we transformed a lactam sensing plasmid into *E. coli* DH5α. The plasmid carries a ChnR-Pb transcription factor-promoter pair that drives mCherry expression upon recognition of lactam by the ChnR protein. We encapsulated the biosensor strain in the 1% agarose core of compartments. 2% agarose gel was used as the shell protective layer. The reconstituted format of lactam biosensors exhibited excellent colony formation and lactam sensing behavior (Fig. 5b). The compartmentalized sensors responded reasonably linearly between 5 mM and 100 mM valerolactam. These results confirm that the ConCEiV core-shell architecture supports the long-term growth and functioning of GMMs.

**Figure 5.**
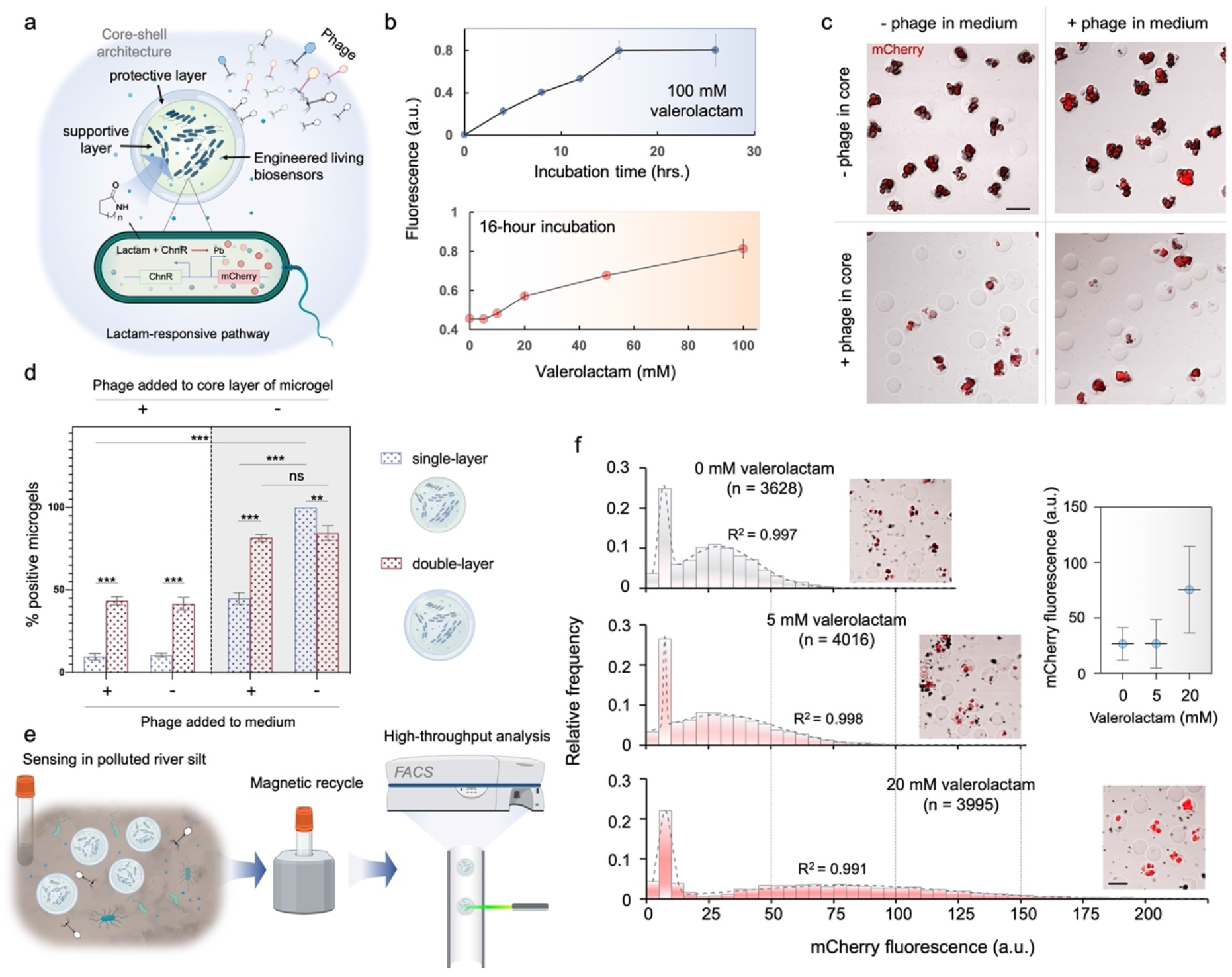
ConCEiV-based living biosensors. (a) Schematic of the compartmentalized biosensor. An engineered *E. coli* strain is encapsulated in the cores (1% agarose) of layered compartments for cell growth and functioning. The strain senses an industrial chemical pollutant – lactam through a constructed lactam-mCherry pathway. A dense outer layer (2% agarose) protects the encapsulated biosensor strain from environmental impacts such as phage attack. (b) Characterization of the compartmentalized biosensors. Top: fluorescence of individual biosensor compartments (n ≥ 50) versus incubation time. The compartments were cultured in medium containing 100 mM valerolactam. Bottom: dose-response relation of the compartmentalized biosensors (n ≥ 50). (c) Optical micrographs showing growth condition of biosensor strain encapsulated in core-shell compartments in presence of phage. The compartments were incubated in medium containing 5 mM valerolactam for 16 hours. The bacteria colonies are labeled in red as they express mCherry fluorescent protein. (d) Comparison of biosensor strain growth in single-layer agarose (1%) beads and in double-layer compartments in presence of phage. Microgels having grown colonies were considered positive. Error bars denote standard deviations of parallel experiments (n = 3). Statistical analysis: unpaired two-tailed t-test. (e) Workflow of sensing, retrieval and analysis of the compartmentalized biosensors in an artificial river silt environment polluted by lactam. (f) Fluorescence profile of compartments after sensing varying levels of valerolactam in polluted river silt. Dashed curves denote dual-peak Gaussian fitting. The micrographs display representative compartments at relevant biosensing conditions. The inset scatter plot shows the mean and standard error of corresponding sensing results. Scale bars: 100 μm.

### Compartmentalized biosensors resist lytic phages

We further investigated whether the layered structure of the compartments could provide adequate protection against phage attacks. Wild DH5α lytic phages were enriched from swine wastewater samples collected at a farm in rural Shanghai. A plaque assay confirmed the phage’s lytic activity on the engineered biosensor strain (Supplementary Fig. 14). We then added the phages to the culture medium and/or to the core of the compartments and examined the cell growth. As depicted in Fig. 5c, colonies formed in most of the compartments with the absence of phages. When the phages were directly added to the core of compartments, the density of colonies dropped significantly, further confirming the phage’s lytic activity. By contrast, the biosensor colonies formed as usual when the phages were only present in the medium. To further confirm the protective effect empowered by the layered ConCEiV architecture, we compared the biosensor growth in bare single-layer (1% agarose, Supplementary Fig. 15) hydrogel beads versus in the layered ConCEiV system (Fig. 5d). As seen, both systems functioned comparably in the absence of phage. With the presence of phages in the medium, however, colonies were found in less than half of the single-layer beads, whereas the difference was negligible for the layered compartments. Even when phages were present in the core gel, over 40% of the compartments were found positive. By contrast, the positive rate of single-layer beads was less than 10%. This might suggest a bidirectional protective effect of the ConCEiV system that it not only resists the phages from outside but also prevents the leakage of phages from inside, thus limiting the spread rate of phages between the compartments. Overall, these results confirm the excellent resistivity of our ConCEiV-based biosensors against predators from the environment, paving the way for their practical usage in the field.

### Long-lasting functioning and retrieval of biosensors in complex environments

To further explore the reconstituted biosensors to survive in ill-defined surroundings, record and memorize desired information, and be retrieved for reading the stored information, we designed an experiment for sensing in a lactam-polluted complex environment (Fig. 5e). In the experiment, the biosensor compartments were put in artificial river silt samples polluted by certain levels of valerolactam. After incubation, they were magnetically purified from the river silts and then subjected to flow cytometry to extract their stored fluorescence information. Antibiotic experiments reveal the competition between wild microbes and the introduced GMMs (Supplementary Fig. 16). The biosensor growth was found markedly slowed down without applying strain-specific antibiotics, yet a duration of 32 hours was found to be sufficient for the biosensors to grow and function. Fig. 5f displays the FACS results of biosensors’ mCherry fluorescence distribution at various pollutant levels. Gaussian fitting was performed on the data, showing that each distribution diagram features two distinct peaks. The first peaks fixed at lower fluorescence (5-10 a.u.) were considered to represent empty compartments or debris. Therefore, we discarded them and derived the fluorescence profile from the second peaks as the biosensor output (Fig. 5f, inset). Clearly, the compartmentalized biosensors successfully recorded corresponding valerolactam levels even in such a complex environment. Notably, the recorded results are consistent with biosensor performance in a culture medium (Fig. 5b), suggesting that re-calibration may not be required when the biosensors are adopted to a strange environment. The above results demonstrate a complete round of living cell-based biosensing in a close-to-reality setting.

## Discussion

A significant advance of ConCEiV is the introduction of configurable semi-permeability to compartmentalized reactors. Realizing such a capability is a daunting task in other established systems like microdroplets. Choosing hydrogel compartments allows direct inheritance of the vast repertoire of composition-property relationships from decades of hydrogel polymer studies. Applying the classic agarose and polyacrylamide hydrogel systems, ConCEiV demonstrates compatibility with several *in vitro* biochemical reaction classes: nucleic acid amplification, TX-TL, and cellular growth. The porous barrier mimics the passive transport phenomena in biology and allows replenishable and sustained biological reactions in the parallel compartments. Other functionalization capabilities of the system such as magnetization improve handling convenience. Further combination of multiple sequential biochemical steps will allow the reconstitution of even more diverse synthetic molecular and cellular systems. The micrometer-sized compartments are readily compatible with commercial analytical tools such as flow cytometers and digital PCR readers. This offers the ease of retrieving stored information from thousands of compartments as independent synthetic biology units.

While our strategy possesses notable advantages, the current manufacturing scheme might impose several limitations. First, the compartment could occasionally break, releasing biological materials in the cavity to the outer environment, ending with the competition for reaction substrates. Second, unlike liposome or cellular membrane, where the core and the outer environment could maintain different liquid compositions, the permeable nature of the compartment precluded the establishment of chemical gradient, as the case in some biological active transport processes. Besides, the tuning of selective permeability solely relies on the adjustment of hydrogel composition during the encapsulation step. It would be favorable if the permeability could be controlled by more active means after the compartment formation. This demand can be potentially met by incorporating temperature- or photo-responsive hydrogel materials^48^.

Owing to its highly configurable nature, the ConCEiV system holds great potential for future synthetic biology applications. For instance, it is possible to encapsulate a large diversity of individual mutants, while performing dedicated reactions separately for directed evolution. As a physical barrier, the compartment could offer protection for the encapsulated living organisms in a native environment, such as human gut^49^, while allowing nutrient exchange with the surroundings. The development of such tools could be helpful to *in vivo* microbiome studies of unculturable bacteria and living bacteria-based therapeutics^36, 50^.

## Acknowledgements

The authors would like to thank Dr. Jinhong Qin for the assistance in bacteriophage experiments. The Confocal Microscopy was conducted at the Analytical Instrumentation Center, and SEM characterization was performed at the Electron Microscopy Center at the School of Physical Science and Technology (SPST), ShanghaiTech University. The fabrication of the microfluidic devices was partially carried out in the Soft Matter Nanofabrication Laboratory of SPST.

## Author Contributions

Y. Liu, J. Zhang, and J. Li directed the research; Y. Liu, J. Zhang, and J. Keasling conceptualize the study. Y. Liu, J. Zhang, L. Li, and R. Zhang designed the experiments. L. Li and R. Zhang carried out the experiments. X. Tian and F. Ba assisted in the cell-free protein synthesis experiments. B. Pu, C. Ma, T. Li, C. Xiong, and Y. Shi contributed to the biosensing experiments. Y. Liu, L. Li, R. Zhang, J. Li, J. Zhang, and J. Keasling analyzed the data and discussed the results. Y. Liu, J. Zhang, and L. Li wrote the manuscript with help from all authors.

## Data and materials availability

All data needed to evaluate the conclusions in the paper are present in the paper and/or the Supplementary Materials. Additional data related to this paper are available from authors upon reasonable request.

## References

1. Szostak, J.W., Bartel, D.P. & Luisi, P.L. Synthesizing life. Nature 409, 387–390 (2001).

2. Rasmussen, S. et al. Transitions from nonliving to living matter. Science 303, 963–965 (2004).

3. Mann, S. The Origins of Life: Old Problems, New Chemistries. Angew Chem Int Edit 52, 155–162 (2013).

4. Ichihashi, N. et al. Darwinian evolution in a translation-coupled RNA replication system within a cell-like compartment. Nat Commun 4 (2013).

5. Thiry, M. & Lafontaine, D.L.J. Birth of a nucleolus: the evolution of nucleolar compartments. Trends Cell Biol 15, 194–199 (2005).

6. Gorlich, D. & Kutay, U. Transport between the cell nucleus and the cytoplasm. Annu Rev Cell Dev Bi 15, 607–660 (1999).

7. Gibcus, J.H. & Dekker, J. The Hierarchy of the 3D Genome. Mol Cell 49, 773–782 (2013).

8. Quinodoz, S.A. et al. RNA promotes the formation of spatial compartments in the nucleus. Cell 184, 5775–5790 e5730 (2021).

9. Pollock, C. & Huang, S. The Perinucleolar Compartment. Csh Perspect Biol 2 (2010).

10. Cremer, T. et al. The Interchromatin Compartment Participates in the Structural and Functional Organization of the Cell Nucleus. Bioessays 42 (2020).

11. Strange, R.E. Effect of Magnesium on Permeability Control in Chilled Bacteria. Nature 203, 1304–1305 (1964).

12. Masi, M., Refregiers, M., Pos, K.M. & Pages, J.M. Mechanisms of envelope permeability and antibiotic influx and efflux in Gram-negative bacteria. Nat Microbiol 2, 17001 (2017).

13. Dam, S., Pages, J.M. & Masi, M. Stress responses, outer membrane permeability control and antimicrobial resistance in Enterobacteriaceae. Microbiology (Reading) 164, 260–267 (2018).

14. Weiss, M. et al. Sequential bottom-up assembly of mechanically stabilized synthetic cells by microfluidics. Nat Mater 17, 89–96 (2018).

15. Joesaar, A. et al. DNA-based communication in populations of synthetic protocells. Nat Nanotechnol 14, 369–+ (2019).

16. Beneyton, T. et al. Out-of-equilibrium microcompartments for the bottom-up integration of metabolic functions. Nat Commun 9, 2391 (2018).

17. Prakadan, S.M., Shalek, A.K. & Weitz, D.A. Scaling by shrinking: empowering single-cell ‘omics’ with microfluidic devices. Nat Rev Genet 18 (2017).

18. Macosko, E.Z. et al. Highly Parallel Genome-wide Expression Profiling of Individual Cells Using Nanoliter Droplets. Cell 161, 1202–1214 (2015).

19. Mashaghi, S., Abbaspourrad, A., Weitz, D.A. & van Oijen, A.M. Droplet microfluidics: A tool for biology, chemistry and nanotechnology. Trac-Trend Anal Chem 82, 118–125 (2016).

20. Tawfik, D.S. & Griffiths, A.D. Man-made cell-like compartments for molecular evolution. Nat Biotechnol 16, 652–656 (1998).

21. Gruner, P. et al. Controlling molecular transport in minimal emulsions. Nat Commun 7 (2016).

22. Deshpande, S., Caspi, Y., Meijering, A.E.C. & Dekker, C. Octanol-assisted liposome assembly on chip. Nat Commun 7 (2016).

23. Shimanovich, U. et al. Silk micrococoons for protein stabilisation and molecular encapsulation. Nat Commun 8 (2017).

24. Huang, X. et al. Interfacial assembly of protein-polymer nano-conjugates into stimulus-responsive biomimetic protocells. Nat Commun 4 (2013).

25. Niederholtmeyer, H., Chaggan, C. & Devaraj, N.K. Communication and quorum sensing in non-living mimics of eukaryotic cells. Nat Commun 9, 5027 (2018).

26. Noireaux, V. & Libchaber, A. A vesicle bioreactor as a step toward an artificial cell assembly. P Natl Acad Sci USA 101, 17669–17674 (2004).

27. Khalil, A.S. & Collins, J.J. Synthetic biology: applications come of age. Nat Rev Genet 11, 367–379 (2010).

28. Way, J.C., Collins, J.J., Keasling, J.D. & Silver, P.A. Integrating Biological Redesign: Where Synthetic Biology Came From and Where It Needs to Go. Cell 157, 151–161 (2014).

29. Harris, D.C. & Jewett, M.C. Cell-free biology: Exploiting the interface between synthetic biology and synthetic chemistry. Curr Opin Biotech 23, 672–678 (2012).

30. Perez, J.G., Stark, J.C. & Jewett, M.C. Cell-Free Synthetic Biology: Engineering Beyond the Cell. Cold Spring Harb Perspect Biol 8 (2016).

31. Cubillos-Ruiz, A. et al. Engineering living therapeutics with synthetic biology. Nat Rev Drug Discov 20, 941–960 (2021).

32. Tang, T.C. et al. Materials design by synthetic biology. Nat Rev Mater 6, 332–350 (2021).

33. Mukherji, S. & van Oudenaarden, A. Synthetic biology: understanding biological design from synthetic circuits. Nat Rev Genet 10, 859–871 (2009).

34. Wan, X. et al. Cascaded amplifying circuits enable ultrasensitive cellular sensors for toxic metals. Nat Chem Biol 15, 540–548 (2019).

35. Sayler, G.S. & Ripp, S. Field applications of genetically engineered microorganisms for bioremediation processes. Curr Opin Biotech 11, 286–289 (2000).

36. Kotula, J.W. et al. Programmable bacteria detect and record an environmental signal in the mammalian gut. P Natl Acad Sci USA 111, 4838–4843 (2014).

37. Ruxton, G.D., Wilkinson, D.M., Schaefer, H.M. & Sherratt, T.N. Why fruit rots: theoretical support for Janzen’s theory of microbe-macrobe competition. Proc Biol Sci 281, 20133320 (2014).

38. Tang, T.C. et al. Hydrogel-based biocontainment of bacteria for continuous sensing and computation. Nat Chem Biol (2021).

39. Abate, A.R., Rotem, A., Thiele, J. & Weitz, D.A. Efficient encapsulation with plug-triggered drop formation. Phys Rev E 84 (2011).

40. Chrambach, A. & Rodbard, D. Polyacrylamide gel electrophoresis. Science 172, 440–451 (1971).

41. Meagher, R.J. et al. End-labeled free-solution electrophoresis of DNA. Electrophoresis 26, 331– 350 (2005).

42. Li, J., Wang, H., Kwon, Y.C. & Jewett, M.C. Establishing a high yielding streptomyces-based cell-free protein synthesis system. Biotechnol Bioeng 114, 1343–1353 (2017).

43. Latulippe, D.R. & Zydney, A.L. Radius of Gyration of Plasmid DNA Isoforms From Static Light Scattering. Biotechnol Bioeng 107, 134–142 (2010).

44. Noeske, J. et al. High-resolution structure of the Escherichia coli ribosome. Nat Struct Mol Biol 22, 336–U389 (2015).

45. Liu, W.Q., Zhang, L.K., Chen, M.Z. & Li, J. Cell-free protein synthesis: Recent advances in bacterial extract sources and expanded applications. Biochem Eng J 141, 182–189 (2019).

46. Jang, S. et al. Artificial Caprolactam-Specific Riboswitch as an Intracellular Metabolite Sensor. Acs Synth Biol 8, 1276–1283 (2019).

47. Mehta, S., Panchal, P., Butala, B. & Sane, S. Bacillus cereus mediated ε-caprolactam degradation: an initiative for waste water treatment of Nylon-6 production plant. Journal of Bioremediation and Biodegradation 5 (2014).

48. Qiu, Y. & Park, K. Environment-sensitive hydrogels for drug delivery. Adv Drug Deliver Rev 64, 49–60 (2012).

49. Cani, P.D. Human gut microbiome: hopes, threats and promises. Gut 67, 1716–1725 (2018).

50. Ruder, W.C., Lu, T. & Collins, J.J. Synthetic Biology Moving into the Clinic. Science 333, 1248–1252 (2011).

