## Supplementary Materials for "Configurable Compartmentation Enables *In Vitro* Reconstitution of Sustained Synthetic Biology Systems"

#### This document includes:

- Materials and Methods
- DNA and protein transport models
- Supplementary Figures and Tables
- References

### Materials and Methods

#### 1. Strains and chemicals

Chemicals used in this study were purchased from Sigma Aldrich unless otherwise stated. The bacteria strains used in this study are listed in Supplementary Table 1. Lactam-responsive plasmids pBbSLactamC-mCherry were extracted from our previously engineered strain *E. coli* DH10B JZ-439[1]. A new lactam sensing strain, *E. coli* LY-439, was constructed by transforming pBbSLactamC-mCherry into a chemically competent DH5α strain. All the *E. coli* strains used in this work were cultured overnight at 30 °C in Luria-Bertani (LB) medium containing 25 µg/mL chloramphenicol.

| Strains | Relevant characteristics | Reference |
|---|---|---|
| *E. coli* DH10B JZ-439 | Lactam responsive<br>Engineered from DH10B<br>F-mcrA crmrr-hsdRMS-mcrBC) r-hsdRMSmcrBC) and oligonucleotide139 ∆ (ara, leu) 7697 galU galK alrpsL nupG | Life Technologies (Carlsbad, CA) |
| *E. coli* DH5α LY-439 | Lactam responsive<br>Engineered from DH5α<br>F- Φ80lacZΔM15 Δ(lacZYA-argF) U169 recA1 endA1 hsdR17(rk-, mk+) phoA supE44 thi-1 gyrA96 relA1 λ- | Life Technologies (Carlsbad, CA) |

**Supplementary Table 1.** Bacteria strains used in this study.

#### 2. Microfluidic device fabrication

The microfluidic devices (supplementary Fig. 1) were designed using AutoCAD. The layout was printed as dark-field plastic photomasks. The devices were fabricated through polydimethylsiloxane (PDMS)-based soft lithography. First, SU-8 3025 photoresist (Microchem) was spin-coated on a 3-inch silicon wafer at desired thickness (~30 µm) following manufacturer's instructions. After a pre-bake of 20 min at 95 °C, the wafer was covered with photomasks and exposed under a 120 mW UV lamp (M365L2, Thorlabs) for 125 s. The wafer was post-baked for 5 min at 95 °C and then developed in a SU-8 developer solution (MicroChem) for 10 min. The fabricated silicon master was further cleaned with isopropanol and ethanol, and blow-dried with a nitrogen gun. PDMS was prepared by mixing its precursor with curing agent

(SYLGARD 184, Dow Corning) at 10:1 (w:w). After vacuum degassing, the PDMS was poured over the master, followed by a curing process of 4 h at 60 °C. The cured PDMS slabs were peeled off from the master and the inlet/outlet ports were created using a custom-made 0.7 mm hole puncher. With the aid of oxygen plasma treatment, each slab was bonded to a clean glass slide. Finally, the fabricated devices were treated with Aquapel (PPG Industries) to render the channel surfaces hydrophobic before use.

#### 3. Compartment manufacturing and characterization

Manufacturing of the compartments started with generating the core hydrogel beads. The hydrogel pre-solution, solution of *in vitro* systems to be encapsulated and a fluorinated oil (HFE-7500, 3M) containing 2 % (w/w) PEG-PFPE amphiphilic block copolymer surfactant (Zhejiang ThunderBio Innovation Ltd.) was injected into a flow-focusing device (supplementary Fig. 1a) at typical flow rates of 300, 200 and 1000 μL/h, respectively, by syringe pumps (NE-501, new era). The flowrate ratio between the aqueous and oil phase could be adjusted between 1:2 to 1:10 to tune the core gel size. After gelling, the emulsion was broken by adding equal volume of 20% (v/v) 1H,1H,2H,2H-perfluoro-1-octanol (Aladdin, catalog no. H157208) in HFE-7500. The excessive oil layer was then discarded. The microgels were further washed by 0.1% (v/v) Triton X-100 in 1X Tris-HCl EDTA (TE) buffer and 0.1% (v/v) Tween 20 (Bioss, catalog no. C-0086) in TE buffer. To form the layered architecture (supplementary Fig. 2a), the core gel beads, pre-solution of the shell layer and HFE-7500 carrying oil were pumped into a dual-junction flow focusing device (supplementary Fig. 1b) at typical flow rates of 100, 300 and 700 μL/h, respectively. The flowrate of shell pre-solution was adjusted between 50 to 300 μL/h to tune the shell thickness. Gelling and de-emulsification steps were then performed as described above. The composition, gelling and re-dissolving condition for the hydrogels are detailed in supplementary Table 2.

| | **Hydrogel** | **Composition (in $ddH_2O$)** | **Gelling condition** | **Dissolving condition** |
|---|---|---|---|---|
| **Core hydrogel** | Polyacrylamide (PAAm)- N,N'- | 6% (w/v) acrylamide, | Add 0.4% (v/v) N,N,N',N'-Tetramethylethylenediamine | 1mM Dithiothreitol |

| | | | | |
|---|---|---|---|---|
| | bis(acryloyl)cystamine (BAC) | 0.392% (w/v) BAC, 0.6% (w/v) ammonium persulfate (APS) | (TEMED) to HFE 7500. Incubate for 30 min | in buffer, 180 s |
| | Agarose | 1% (w/v) ultra-low melting temperature agarose | Set on ice for 10 min | 45 °C, 10 min |
| **Shell hydrogel** | Bis-PAAm (3) | 6% (w/v) acrylamide, 0.6% (w/v) N'-methylene bis(acrylamide) (Bis), 0.45% (w/v) APS | Add 0.4% (v/v) TEMED to HFE 7500. Incubate for 30 min | |
| | Agarose | 2% (w/v) ultra-low melting temperature agarose | Set on ice for 10 min | |

**Supplementary Table 2.** Composition, gelling and dissolving conditions for layered compartments displayed in Fig. 2

The re-dissolving of core gel of compartments was first tested on bare PAAm-BAC core gel beads (Supplementary Fig. 3). The beads were successfully dissolved in 90 s as confirmed under bright field (Supplementary Fig. 3a). The dissolving process were further confirmed with fluorescent microscopy using Cy3-DNA labeled beads (Supplementary Fig. 3b). The labeled beads were manufactured by adding 100 nM acrydite-modified DNA primers (5'-acrydite-TAG CTT ATC AGA CTG ATG TTG A-3') to the pre-gel solution and co-incubating the beads with 1 μM cy3-labeled complimentary sequence. To confirm the all-liquid cavity formed after core dissolving, the Cy3-DNA label core beads were further manufactured to core-shell architecture (Fig. 2d). The shell of the core-shell architecture contained 1mg/ml of FITC-dextran (1000 kDa) for fluorescent visualization. Dissolving the core of the core-shell architecture features a slower dynamic of 180 s (Supplementary Fig. 4). The dissipated Cy3 fluorescence of the compartments (Fig. 2d) verifies that the Cy3-DNA molecules transported to the outer environment, confirming the complete dissolving of core hydrogel polymer network. The fluorescent microscopy was performed with a confocal microscope (Olympus LS FV3000). The compartments were magnetized by co-encapsulating 50 nm $SiO_2$- $Fe_3O_4$ magnetic nano particles (Daojin Tech. Co.) in the shell layer. The magnetism of compartments was confirmed by a magnetic cleaning experiment (Supplementary Fig. 5).

To obtain the pore size profile of the compartments, three Bis-PAAm hydrogel samples were examined by scanning electron microscopy (SEM). The composition of the samples is detailed in Supplementary Table 3. The hydrogel samples were first dehydrated in an ascending series of ethanol aqueous solutions (25%, 50%, 75%, 90%, and 100% /v). The dehydrated gels were then further dried through a $CO_2$ critical point drying (CPD) procedure[2]. The CPD-dried samples were carefully cut to expose their internal porous structures and sputtered with gold (20 nm). SEM micrographs were taken using a field emission high resolution scanning electron microscopy (JSM-IT500HR/LA) at an operating voltage of 5 kV. The SEM images are shown in Supplementary Fig. 6.

| | Sample 1 | Sample 2 | Sample 3 |
|---|---|---|---|
| Acrylamide | 9% | 9% | 6% |
| Bis | 0.24% | 0.45% | 0.6% |
| APS | 0.45% | 0.45% | 0.45% |
| TEMED | 0.4% | 0.4% | 0.4% |

**Supplementary Table 3.** Composition of three Bis-PAAm hydrogel samples subject to SEM.

#### 4. Compartmentalized PCR

The compartmentalized PCR reaction contains 19 µL Bis-PAAm compartments containing plasmid pFB2[3] (10 nM final concentration), 25 µL 2X Phanta Max Buffer (Vazyme, catalog no. P505-d1), 1 µL 10mM dNTP Mix (Vazyme, catalog no. P505-d1), 1 µL 10 µM forward primer, 1 µL 10 µM reverse primer and 1 µL Phanta Max Super-Fidelity DNA Polymerase (Vazyme, catalog no. P505-d1). The reaction was performed in a BioRad T100 thermal cycler using the following protocol: 95 °C for 3 min, followed by 35 cycles of 95 °C for 15 s, 58 °C for 15 s, 72 °C for 3 min, a final extension at 72 °C for 5 min and holding at 4 °C. After PCR, the compartments were magnetically cleaned to removed excessive DNA product in the buffer and resuspended in 1X Tris-HCl buffer (Supplementary Fig. 7). Gel electrophoresis (1% TAE agarose gel) was performed on the compartments to determine that desired fragments were amplified (Supplementary Fig. 8). The samples were finally examined under the FAM channel of a digital droplet PCR reader (Nebula, Zhejiang ThunderBio Innovation Ltd.) to extract the fluorescence of individual compartments (Supplementary Fig. 9). The primer pairs targeting 150-1187 bp fragments on the plasmid were designed using an online tool PRIMER3. All the primers were purchased from Genewiz Co. Ltd.. Their sequences are listed in Supplementary Table 4.

| Index | Forward primer (5'-3') | Reverse primer (5'-3') | Product size |
|---|---|---|---|
| 1 | TGTTTACATCACCGCCGATA | TGATTGTCTGGCAGCAGAAC | 150 bp |
| 2 | CTGGCTGATCACTACCAGCA | GGTACCGTCGACTGCAGAAT | 225 bp |
| 3 | GGGCGAAGAAGTTGTCCATA | TCCGGCCTTTATTCACATTC | 338 bp |
| 4 | GGTAAACTGCCGGTACCTTG | AGCAGAACAGGACCATCACC | 434 bp |
| 5 | CGCATGGTATGGATGAACTG | TTTGAGCGTCAGATTTCGTG | 547 bp |
| 6 | CTCGAGGCTTGGATTCTCAC | TCCGGCCTTTATTCACATTC | 620 bp |
| 7 | CTCGAGGCTTGGATTCTCAC | ATCCCAATGGCATCGTAAAG | 749 bp |
| 8 | TCTGACGCTCAAATCAGTGG | TGTCGGCAGAATGCTTAATG | 866 bp |
| 9 | TGGGCCATAAGCTGGAATAC | AGGCGTGGAATGAGACAAAC | 968 bp |
| 10 | CTGTTGTTTGTCGGTGAACG | GATATCGAGCTCGCTTGGAC | 1187 bp |

**Supplementary Table 4.** Primer sequences used in the compartmentalized PCR experiments.

#### 5. Compartmentalized cell-free protein synthesis (CFPS)

PAAm-Bis compartments with liquid cavities (cavity diameter: ~60 µm, shell thickness: ~15 µm) encapsulating sfGFP were manufactured as described above. All the compartments encapsulated 13.3 µg/mL plasmid; for CFPS, equal volume of crude cell extract was co-flowed with the pre-gel solution; an extra dose of 2.5-unit T7 RNA polymerase (Invitrogen, 18033019) was added in some cases. To obtain crude cell extract, *E. coli* BL21 Star (DE3) strain was cultured in 2×YTPG medium (10 g/L yeast extract, 16 g/L tryptone, 5 g/L NaCl, 7 g/L $K_2HPO_4$, 3 g $KH_2PO_4$, and 18 g/L glucose, pH 7.2) in 2.5 L baffled Ultra Yield™ flasks (Thomson Instrument Company). After inoculation of 20 mL overnight cultures, cells were incubated in the shaker at 220 rpm 34 °C (initial $OD_{600}$ of 0.05-0.1). When $OD_{600}$ reached 0.6–0.8, isopropyl-β-D-thiogalactopyranoside (IPTG) was added to a final concentration of 1 mM to induce T7 RNA polymerase expression. When $OD_{600}$ reached ~3.0, cells were harvested by centrifugation at 4500 rpm, 4 °C for 15 min. After that, cell pellets were washed three times with cold S30 Buffer (10 mM Tris-acetate, 14 mM magnesium acetate, 60 mM potassium acetate, and 2 mM DTT), resuspended (1 mL per gram of wet cell mass) and lysed by sonication (Q-sonica, 10 s on/off, 50% amplitude). The lysate was then centrifuged twice at 12, 000 g, 4 °C for 10 min. The resulting supernatant was flash frozen in liquid nitrogen and stored at -80 °C for further use.

A typical 45 µL compartmentalized CFPS assay includes 12 mM magnesium L-glutamate, 10 mM ammonium L-glutamate, 130 mM potassium L-glutamate, 1.2 mM ATP, 0.85 mM each of GTP, UTP, and CTP, 34 µg/mL folinic acid, 170 µg/mL of *E. coli* tRNA mixture, 2 mM each of 20 standard amino acids, 0.33 mM nicotinamide adenine dinucleotide (NAD), 0.27 mM coenzyme A (CoA), 1.5 mM spermidine, 1 mM putrescine, 4 mM sodium oxalate, 33 mM phosphoenolpyruvate (PEP), 27% (v/v) of cell extract and 20 µL of plasmid encapsulating compartments. All the reactions were incubated at 30 °C for 6 h.

To monitor mRNA levels during the process, the assays were terminated at the stated durations and purified with Eastep Super Total RNA Extraction Kit (Promega) to extract the mRNA. Gel electrophoresis was performed to confirm the success of mRNA extraction (data not shown). Immediately after extraction, mRNA samples were quantified by reverse transcription-PCR (RT-PCR) using a One-Step PrimeScript™ RT-PCR Kit (TaKaRa, Code No. RR064A). The RT-PCR protocol was 42 °C for 5 min, 95 °C for 10 s, 35 cycles of (95 °C for 15 s, 56 °C for 15 s and 72 °C for 1 min), final extension at 72 °C for 5 min and holding at 4 °C. A standard curve was first derived for quantification of mRNA concentrations (Supplementary Fig. 10). For comparison, mRNA levels of a 45 µL bulk CFPS reaction were also monitored.

For multicycle protein synthesis, the magnetic compartments were purified from a 45 µL compartmentalized CFPS assay on a custom-made magnet rack after 6-hour incubation and immediately immersed in a new tube containing the CFPS mixture. The supernatant of each reaction was collected to analyze sfGFP yield. To quantify the absolute mass concentration of sfGFP, a standard curve describing the sfGFP mass concentration versus the fluorescence intensity was derived (Supplementary Fig. 11). To obtain the curve, His-tagged sfGFP expressed *in vivo* was purified by a nickel column and then diluted into 45 μL CFPS mixture (excluding the sfGFP plasmid) at varying concentrations. The fluorescence of known sfGFP concentrations was read by a plate reader (BioTek synergy H1). Supplementary Fig. 12 shows the sfGFP yield along a 6-hour CFPS reaction with only cell extract in the compartment cavity and with extra 2.5 U of purified T7 RNA polymerase. The sequence structure of the sfGFP plasmid used in this study is illustrated in Supplementary Fig. 13.

**6. Compartmentalized living bacteria-based biosensors**

To constructed the compartmentalized living bacteria-based biosensors, the core-shell architecture compartments were used. The compartments (50 μm core diameter, 10 μm shell thickness) were composed by 1% ultra-low agarose gel as core and 2% ultra-low agarose gel (containing MNPs) as protective shell. The cores encapsulated *E. coli* LY-439 cells at an average density of 5 cells/compartment. To characterize lactam sensing performance of the compartmentalized sensors, the compartments were cultured in LB medium containing 25 μg/mL chloramphenicol, the stated concentrations of valerolactam (Fig. 5b) and 0.002% (w/w) arabinose (BIOSYNTH, catalog no. A-8240). The compartments were imaged at the stated time durations under a microscope (Olympus LS FV3000) and the fluorescence profile of compartments were analyzed by the software ImageJ (http://rsb.info.nih.gov/ij).

Wild bacteriophages were enriched and purified from swine wastewater samples collected from a farm in Jinshan district of Shanghai, China following an established protocol[4]. A phage lysis experiment was performed to confirm the lytic activity of the purified phages on strain *E. coli* LY-439 (Supplementary Fig. 14). The purified bacteriophage solution was stored at 4 °C or mixed with 20 % glycerin and stored at -80 °C for further use. To evaluate the protective effect of the compartments on phage attack, the purified bacteriophage solution was spiked in the culture medium at ~1:1000 v/v or spiked in the core pre-gel solution at ~1:100 v/v. All the cultures were incubated at 30 °C for 16 h (unless otherwise stated). After incubation, the core-shell compartments or single-layer agarose beads were washed by magnetic separation, resuspended in 1X PBS buffer and imaged by fluorescent microscopy (Fig. 5c and Supplementary Fig. 15). Colony formation within the compartments was manually analyzed. The compartments/beads having visible bacteria colonies were considered positive.

To examine the behavior of compartmentalized biosensors in complex environments, we collected pond silty water samples from ShanghaiTech University campus. The samples were spiked in with the stated concentrations of valerolactam, 0.002% (w/w) arabinose and 25 μg/mL chloramphenicol (CmR) to mimic polluted river silty water. Before the sensing experiments, the compartmentalized biosensors were cultured in the silt samples (both CmR- and CmR+) and LB medium at 30 °C for stated duration to investigate the growth of encapsulated biosensor strain (Supplementary Fig. 16). Regarding the final biosensing experiments, the compartments were cultured in artificial river silt samples at room temperature for 16 h. The compartments were then magnetically purified and resuspended in 1X PBS buffer. Finally, flow cytometry was performed on the clean compartment samples to analyze fluorescence of individual compartments using a benchtop flow cytometer (Namocell). All data in this work was analyzed and plotted using GraphPad prism (https://www.graphpad.com/) or Microsoft Excel. The Figures were designed with BioRender (https://www.biorender.com/) or Microsoft Powerpoint.

#### DNA and protein transport models

**a) DNA transport model**

One can estimate the approximate radii ($R_G$) of gyration of DNA fragments with the Kratky-Porod equation[5], given as

$$R_G^2 \approx \frac{b_K M b}{6}\left[1 - 3\left(\frac{b_K}{2Mb}\right) + 6\left(\frac{b_K}{2Mb}\right)^2 - 6\left(\frac{b_K}{2Mb}\right)^3\left(1 - e^{-\frac{2Mb}{b_K}}\right)\right] \quad \text{(S1)}$$

where $b_K$ is the polymer Kuhn length. $L_D$ represents the total contour length of polymer, given by

$$L_D = Mb \quad \text{(S2)}$$

Where $M$ is the number of monomers of a polymer molecule and $b$ is monomer size. In our model, we approximate that a DNA molecule is not able to transport across the hydrogel shell when its diameter (double of its $R_G$) is greater than the hydrogel pore size ($D_p$). The confined-to-diffusible transition takes place at a critical situation where

$$D_p = 2R_G \quad \text{(S3)}$$

Combining Eqs. S1-S3, we can derive the relation between a given $D_p$ and the critical number of monomers of a DNA molecule right experiencing the confined-to-diffusible transition ($M_{crit}$), given as

$$D_p = 2\left\{\frac{b_K M_{crit} b}{6}\left[1 - 3\left(\frac{b_K}{2M_{crit}b}\right) + 6\left(\frac{b_K}{2M_{crit}b}\right)^2 - 6\left(\frac{b_K}{2M_{crit}b}\right)^3\left(1 - e^{-\frac{2M_{crit}b}{b_K}}\right)\right]\right\}^{\frac{1}{2}} \quad \text{(S4)}$$

Eq. S4 divides the diffusible regime and confined regime of DNA transport through the compartment shell. For DNA, a typical estimation for $b_K$ and $b$ is that $b_K \approx 100$ nm and $b \approx 0.34$ nm[5].

**b) protein transport model**

In the protein transport model, we consider the gyration radius of a globular protein ($R_{Gp}$) as[6]

$$R_{Gp} = 0.395 \times N^{3/5} + 7.257 \quad \text{(S5)}$$

Where N denotes the total number of amino acids of a protein molecule. Eq. S5 performed excellent estimation of a dataset of ~1000 globular proteins from RCSB protein data bank database (https://www.rcsb.org/)[7]. Similar to the DNA model, we can derive the relation between a given $D_p$ and the critical number of amino acids of a protein molecule right experiencing the confined-to-diffusible transition ($N_{crit}$), given as

$$D_P = 2\left(0.395 \times {N_{crit}}^{3/5} + 7.257\right) \quad \text{(S6)}$$

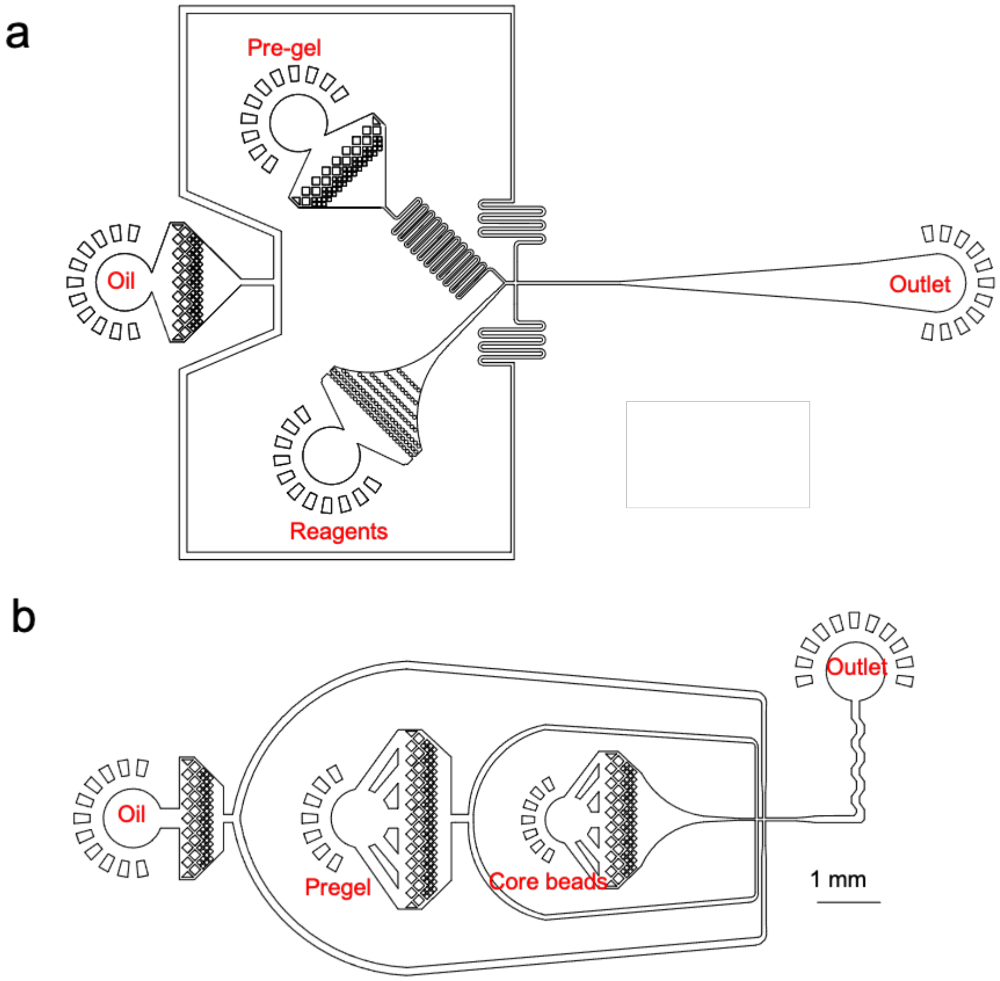


**Supplementary Figure 1.** The layout of microfluidic devices for **a)** generating core hydrogel beads and **b)** core beads reinjection and core-shell compartments manufacturing.

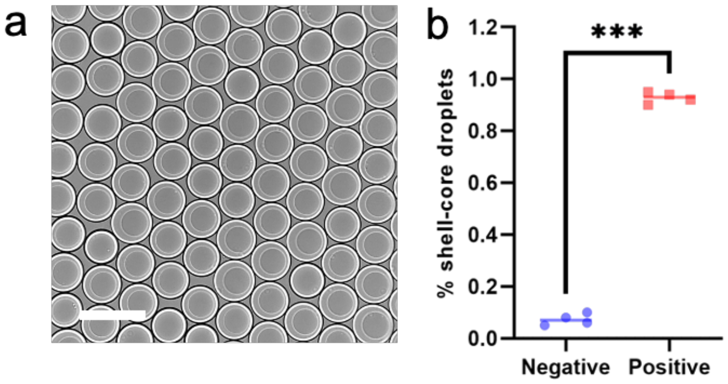

**Supplementary Figure 2.** Generating core-shell microdroplets. **a)** microscopic image of as-generated core-shell droplets. Scale bar: 100 μm. **b)** Statistics of droplets containing core beads (positive) and bare droplets (negative).

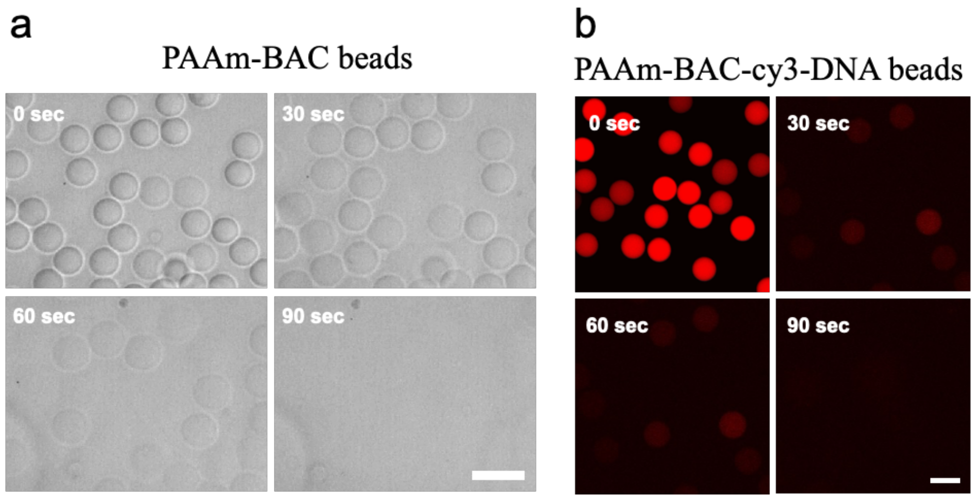


**Supplementary Figure 3.** Re-dissolving core PAAm-BAC beads. **a)** Serial microscopic images of dissolving PAAm-BAC beads with 1 mM DTT. **b)** Serial fluorescence images of dissolving PAAm-BAC-cy3-DNA beads with 1 mM DTT. Scale bars: 50 μm.

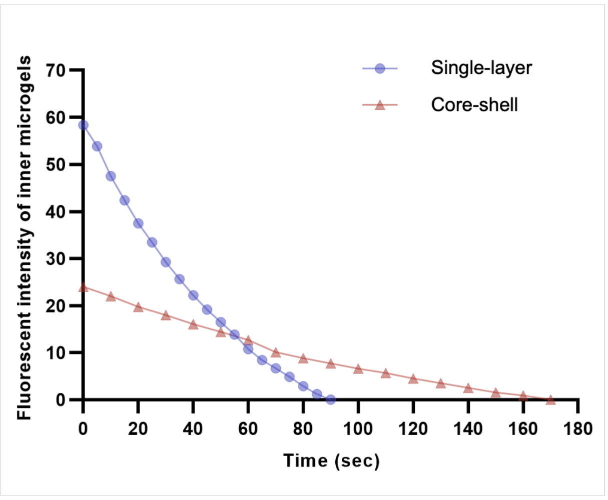


**Supplementary Figure 4.** Comparative analysis of the re-dissolving dynamics of single-layer PAAm-BAC beads and their core-shell format. The dissolving process of the core-shell compartments features slower dynamics due to limited diffusion of DTT.

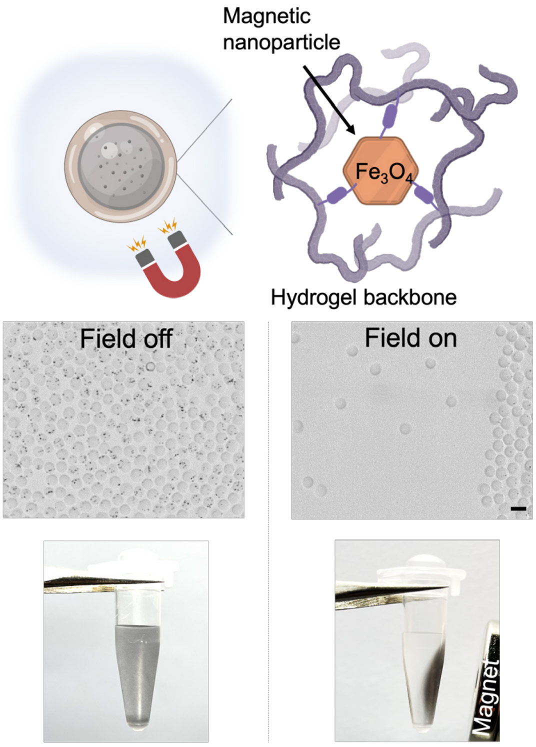


**Supplementary Figure 5.** Magnetizing of the compartments by encapsulating $Fe_3O_4$-$SiO_2$ MNPs in the shell layer. The magnetized compartments can be attracted by a hopper magnet commonly available in biology laboratories, thus enabling convenient magnetic purification and liquid exchange of the compartments. Scale bar: 100 µm.

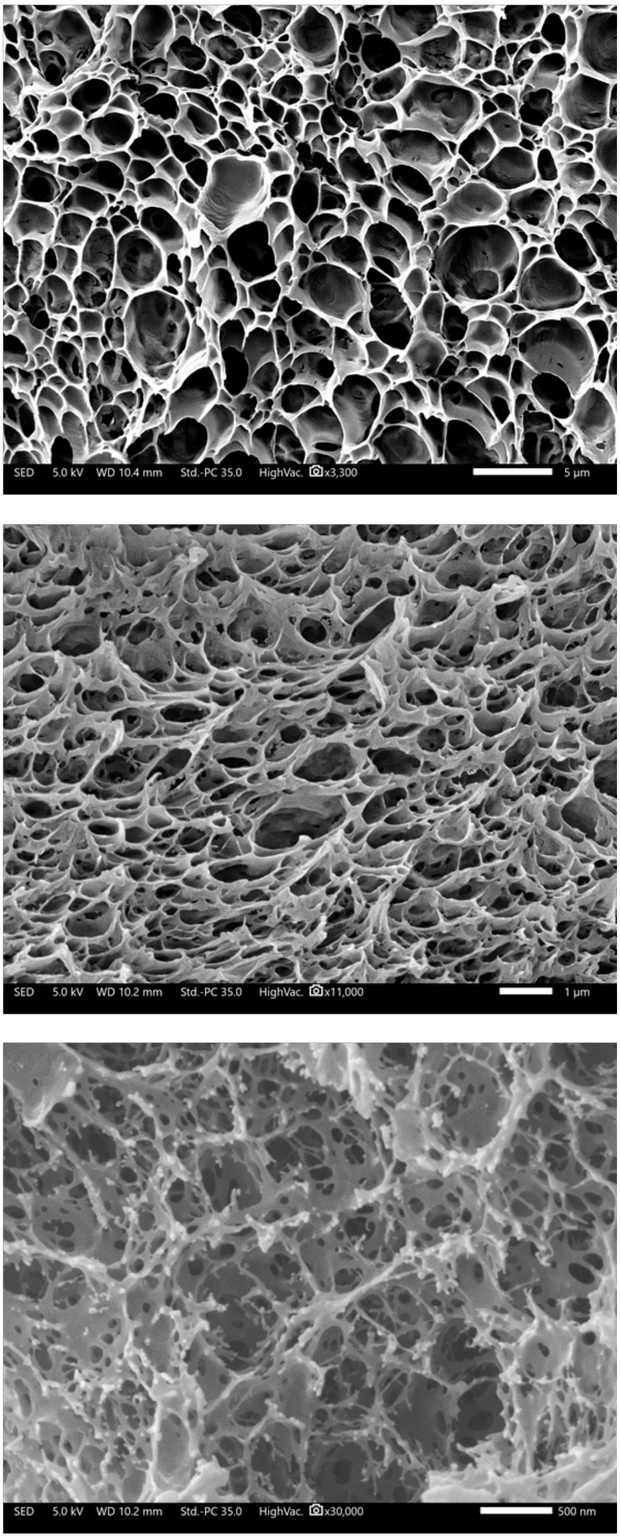


**Supplementary Figure 6.** Representative scanning electron micrographs of Bis-PAAm hydrogels containing **a)** 0.24%, **b)** 0.45% and **c)** 0.6%. The pore sizes of these hydrogel materials were measure as 131±51, 58±13, 31±5 nm for 0.24%, 0.45% and 0.6% Bis-PAAm hydrogel, respectively (n = 20). The composition detail of these hydrogels is described in Supplementary Table 3.

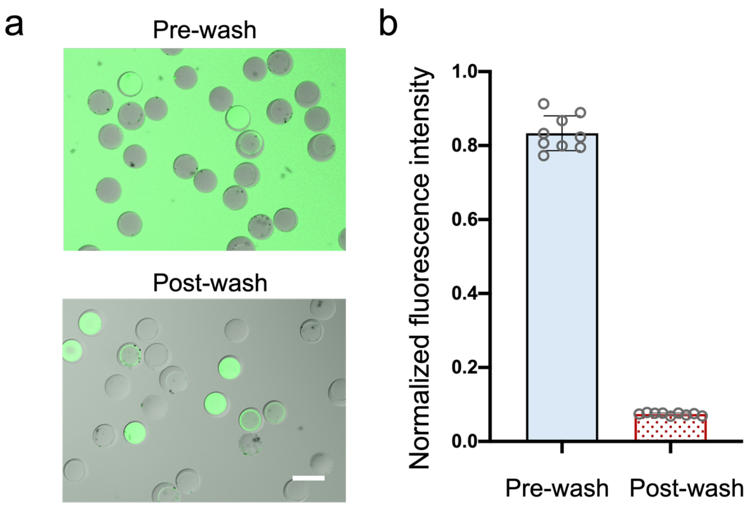


**Supplementary Figure 7.** A compartmentalized PCR assay. **a)** fluorescent images and **b)** background fluorescence intensity of the PCR mixture prior to and after magnetic cleaning. The dsDNA products were labeled by SYBR green dye. Scale bar: 100 µm.

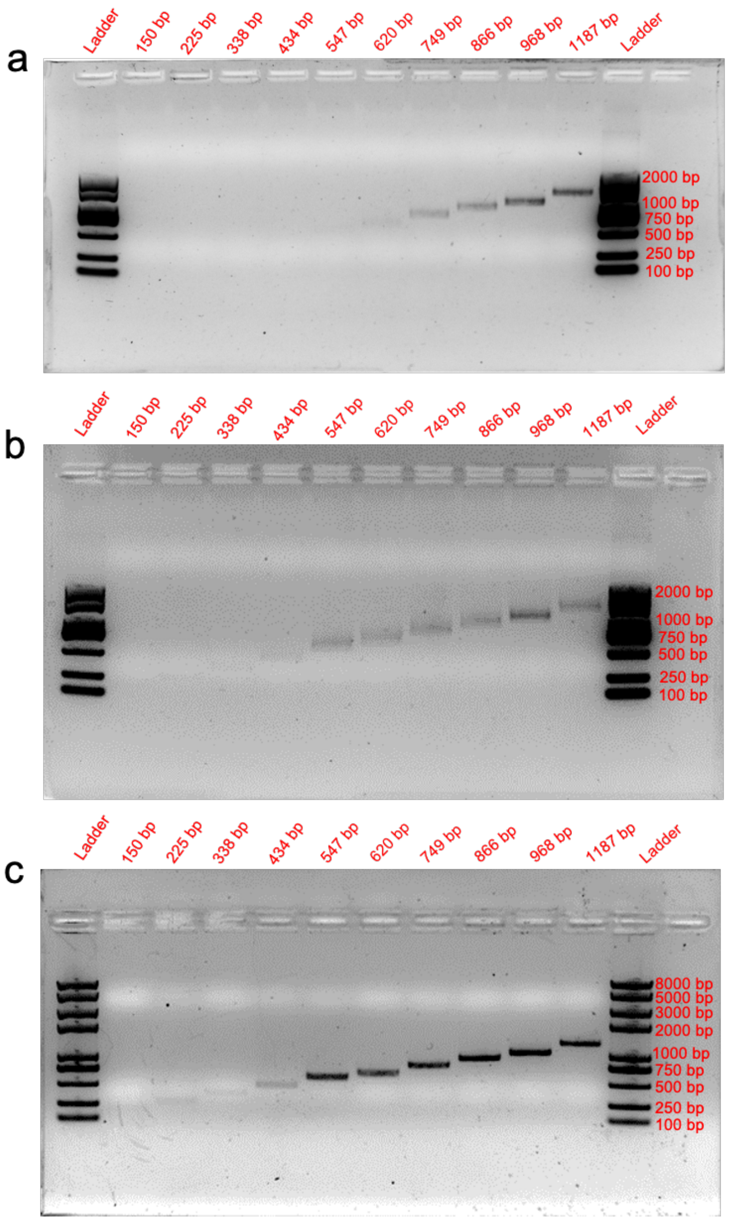


**Supplementary Figure 8.** Gel electrophoresis results of PCR assays performed with **a)** 0.24% **b)** 0.45% and **c)** 0.6% Bis-PAAm core-shell compartments. The expected size of amplicons was 150, 225, 338, 434, 547, 620, 749, 866, 968 and 1187 bp.

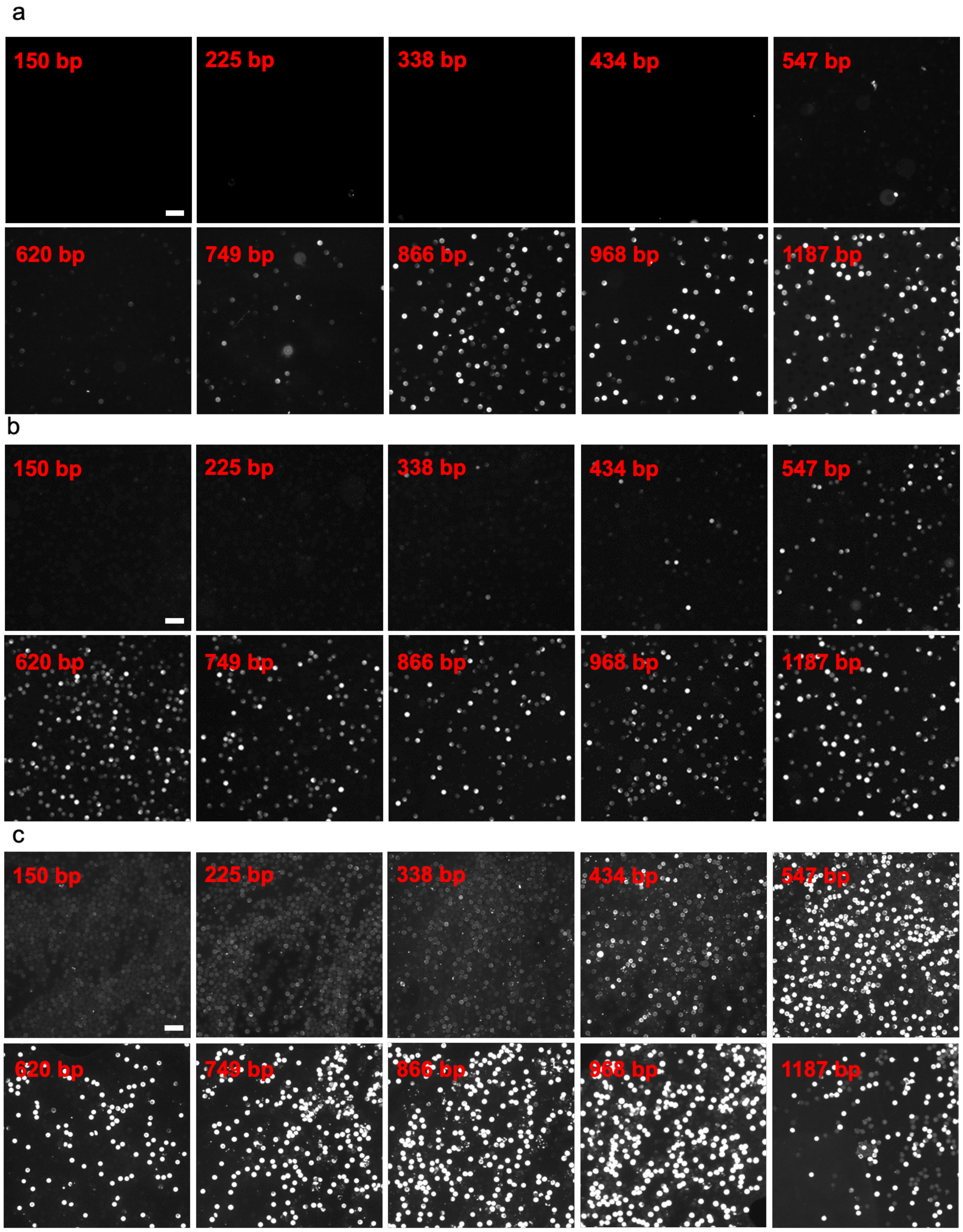


**Supplementary Figure 9.** Fluorescent micrographs of PCR assays performed with **a)** 0.24% **b)** 0.45% and **c)** 0.6% Bis-PAAm core-shell compartments. Before imaging, the samples were magnetically purified to remove the background. Scale bars, 200 µm.

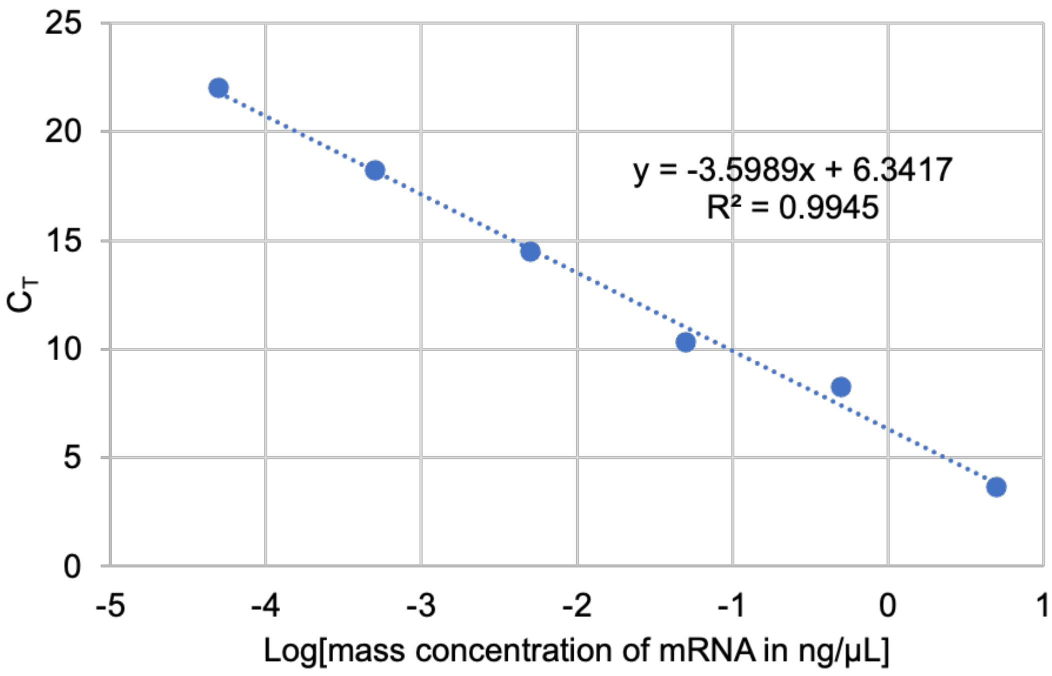


**Supplementary Figure 10.** Standard curve of mRNA mass concentration versus the threshold cycle ($C_T$) of RT-PCR. This curve was used to quantify the total mRNA level during a compartmentalized CFPS reaction of sfGFP.

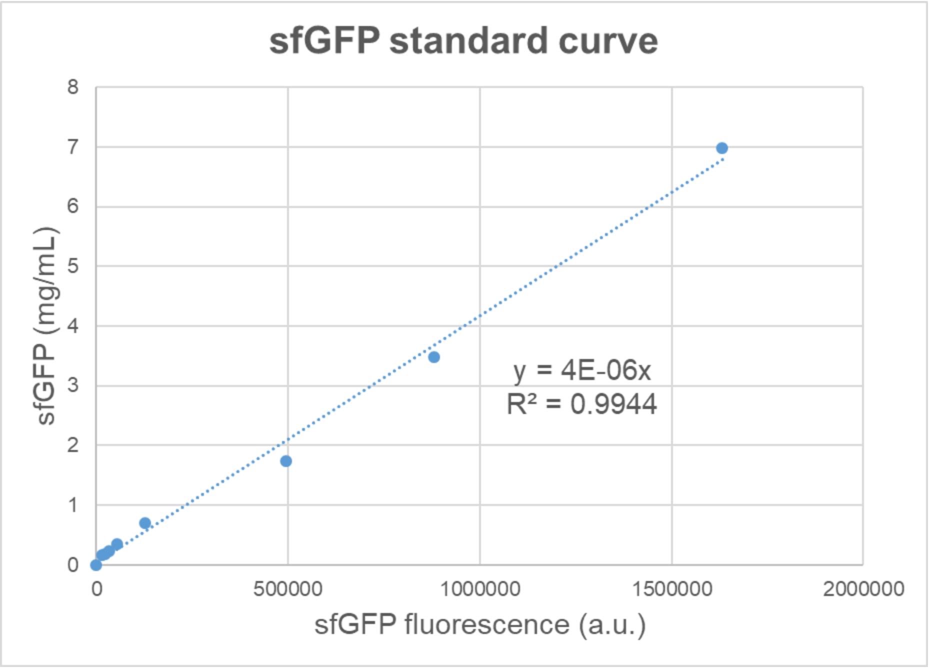


**Supplementary Figure 11.** Standard curve of sfGFP protein concentration versus fluorescence intensity.

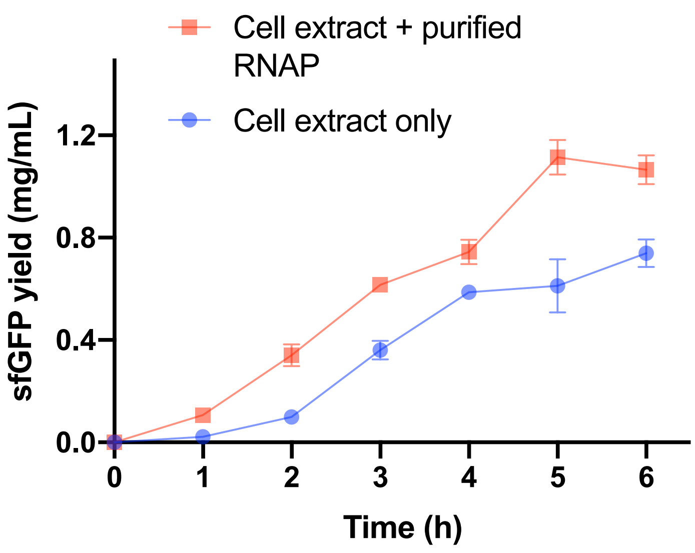


**Supplementary Figure 12.** sfGFP yield plotted over incubation time in compartmentalized CFPS systems with only cell extract (blue) and extra 2.5 U T7 RNAP (red) in the compartment cavity. Both experiments contained sfGFP plasmid in the compartments.

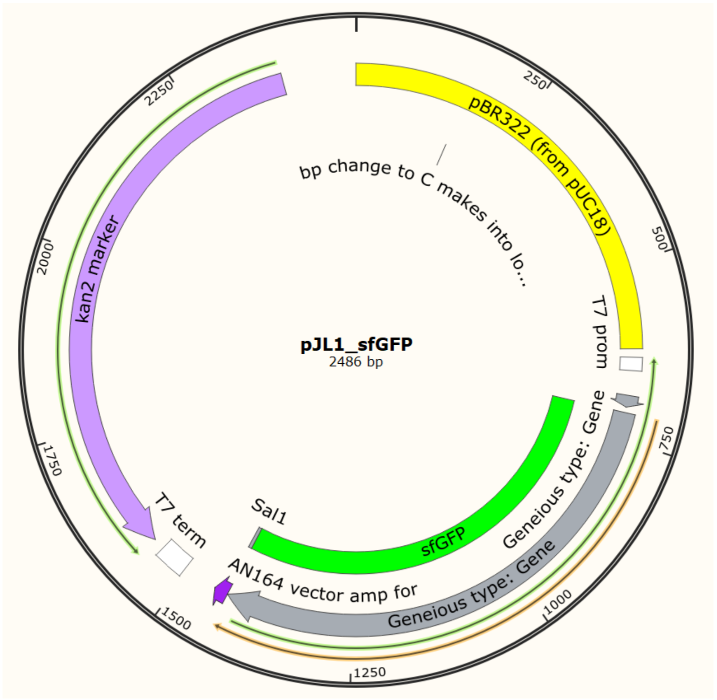


**Supplementary Figure 13.** Map of the sfGFP plasmid used in this work.

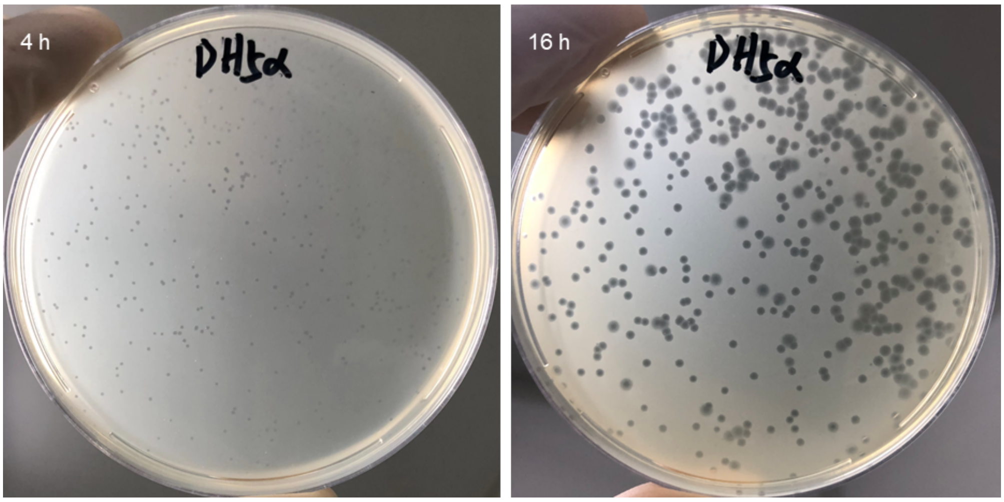


**Supplementary Figure 14.** Bacteriophage lytic activity test. The phage solution was sprayed on an agar plate cultured with *E. coli* LY-439 and after 4 and 16 hours the images were taken. The plaques gradually appearing and expanding on the agar surface confirmed the lytic activity of the wild phages against the *E. coli* strain LY-439.

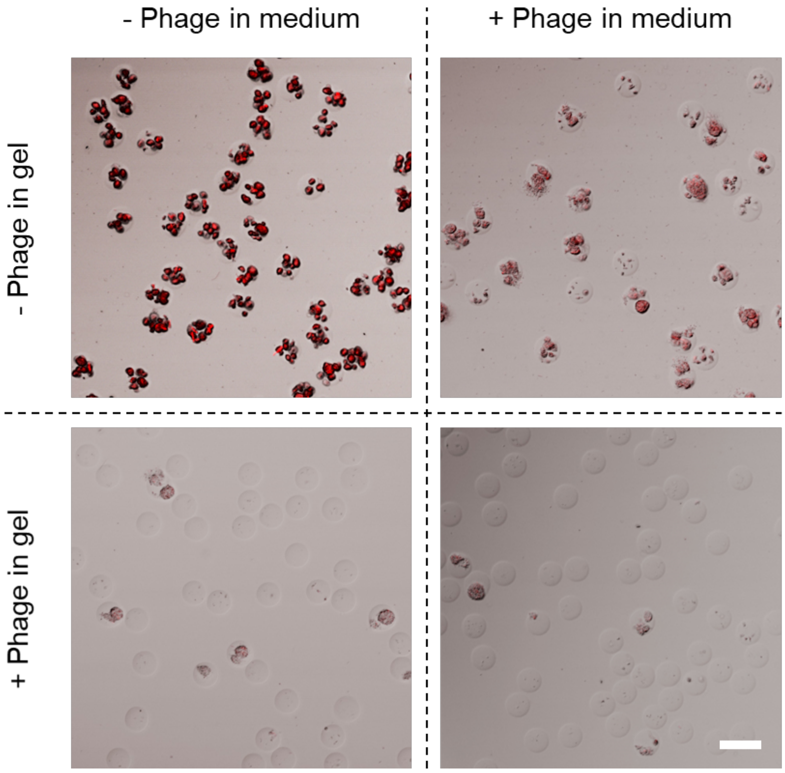


**Supplementary Figure 15.** Phage lysis experiment performed on single-layer agarose beads encapsulating the biosensing bacteria strain. The images display the gel beads after 16-hour culture in LB medium (with 5 mM valerolactam) with the stated phage existence. Scale bar: 100 µm.

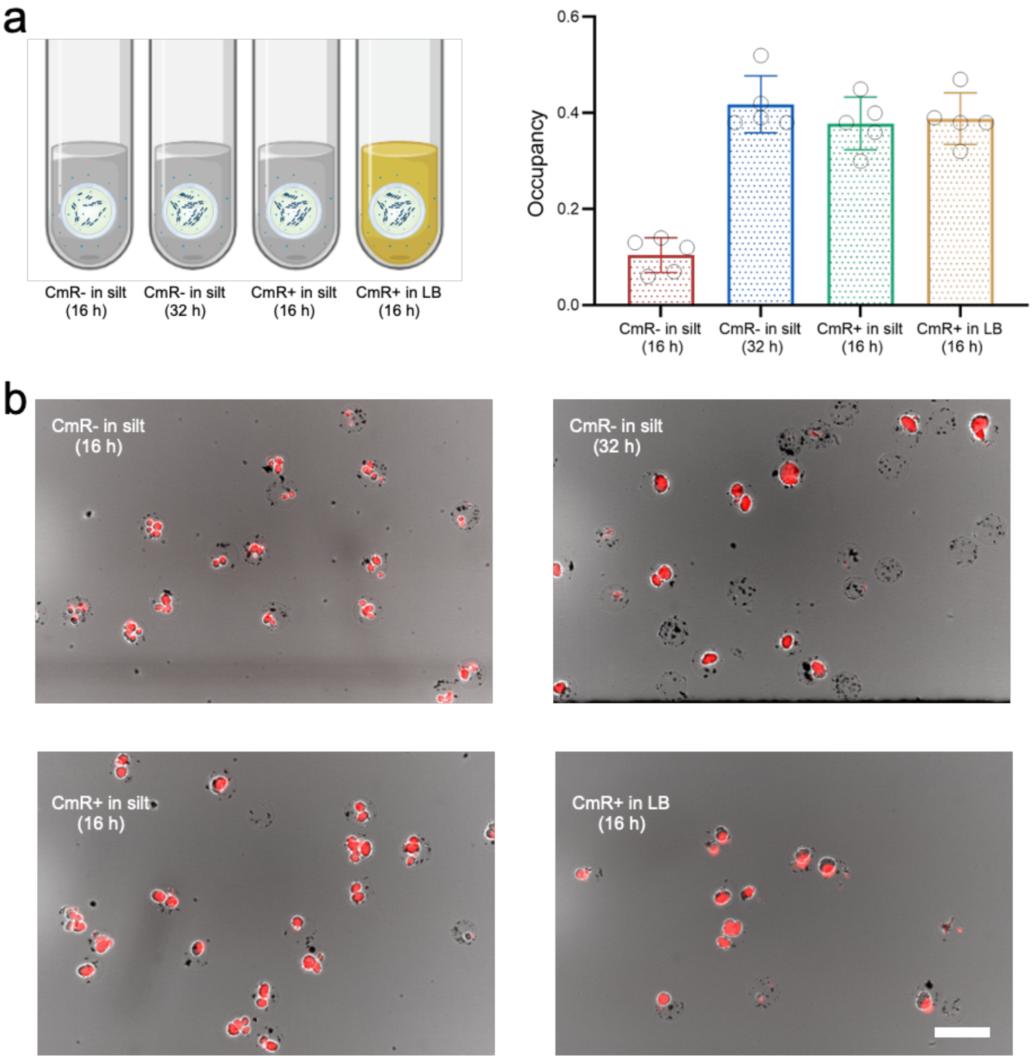


**Supplementary Figure 16.** Growth and function of the compartmentalized biosensors in artificial river silt environments. **a)** the living bacteria-based biosensors were casted into artificial river silt samples under the stated conditions. An LB medium culture experiment was simultaneously conducted for comparison. After the stated duration, compartments were recovered from the samples by magnetic cleaning and the cell growth behavior was analyzed by calculating the occupancy of colony [Area(colony)/area(compartment)]. **b)** microscopic images displaying the biosensor compartments subject to the stated culturing conditions. The bacteria colonies are labeled in red as they express mCherry fluorescent protein.

#### References


1. Zhang, J. et al. Development of a Transcription Factor-Based Lactam Biosensor. *ACS Synth Biol* **6**, 439-445 (2017).
2. Hsu, M.N. et al. Smart Hydrogel Microfluidics for Single-Cell Multiplexed Secretomic Analysis with High Sensitivity. *Small* **14**, e1802918 (2018).
3. Liu, W.S., Zhao, H.Y., Jia, B., Xu, L. & Yan, Y.J. Surface display of active lipase in Saccharomyces cerevisiae using Cwp2 as an anchor protein. *Biotechnol Lett* **32**, 255-260 (2010).
4. Van Twest, R. & Kropinski, A.M. Bacteriophage enrichment from water and soil. *Methods Mol Biol* **501**, 15-21 (2009).
5. Meagher, R.J. et al. End-labeled free-solution electrophoresis of DNA. *Electrophoresis* **26**, 331-350 (2005).
6. Li, X. et al. The repeat region of cortactin is intrinsically disordered in solution. *Sci Rep-Uk* **7** (2017).
7. http://www.scfbio-iitd.res.in/software/proteomics/rg.jsp.
